# *SignalingProfiler* 2.0: a network-based approach to bridge multi-omics data to phenotypic hallmarks

**DOI:** 10.1101/2024.01.25.577229

**Authors:** Veronica Venafra, Francesca Sacco, Livia Perfetto

## Abstract

Unraveling the cellular signaling remodeling upon a perturbation is a fundamental challenge to understand disease mechanisms and to identify potential drug targets. In this pursuit, computational tools that generate mechanistic hypotheses from multi-omics data have invaluable potential. Here we present *SignalingProfiler* 2.0, a multi-step pipeline to systematically derive context-specific signaling models by integrating proteogenomic data with prior knowledge-causal networks. This is a freely accessible and flexible tool that incorporates statistical, footprint-based, and graph algorithms to accelerate the integration and interpretation of multi-omics data. Through benchmarking and rigorous parameter selection on a proof-of-concept study, performed in metformin-treated breast cancer cells, we demonstrate the tool’s ability to generate a hierarchical mechanistic network that recapitulates novel and known drug-perturbed signaling and phenotypic outcomes. In summary, S*ignalingProfiler* 2.0 addresses the emergent need to derive biologically relevant information from complex multi-omics data by extracting interpretable networks.

## Introduction

Intracellular signaling pathways, marked by molecular interactions and post-translational modifications like phosphorylation, mediate the ability of cells to translate signals into observable changes in phenotypic traits. Numerous pathways (e.g., MAPKs, EGFR,…) have been extensively studied and it is now evident that these linear cascades are not isolated entities, but rather components of a large and complex network that impact physiological and pathological processes (Jordan *et al*, 2013). To understand the intricate nature of such a human naïve signaling network it is crucial to grasp the cross-talk among diverse signaling cascades and elucidate how they collectively impact key cellular phenotypes.

In this scenario, the recent tremendous technological advances have enabled the cost-effective generation of large-scale-omics datasets, providing a systematic description of different regulatory layers (e.g., DNA, RNA, and protein levels) in various pathophysiological conditions. The simultaneous exploration of different omics layers (the so-called, “trans-omics analysis”) is indeed gaining popularity (Wu *et al*, 2021b; Terakawa *et al*, 2022; Wu *et al*, 2021a) to obtain a holistic picture of the cell state (Mohammadi-Shemirani *et al*, 2023). However, extracting biological information from such complex omics data remains a major challenge and demands computational interventions.

Among the different methods developed (Reimand *et al*, 2019; Cantini *et al*, 2017), footprint-based techniques (Dugourd & Saez-Rodriguez, 2019; Schubert *et al*, 2018) generate lists of kinases and transcription factors characterized by an activity score derived from the phosphorylation or expression level of their known targets (Mercatelli *et al*, 2020; Beekhof *et al*, 2019; Mischnik *et al*, 2016; Badia-I-Mompel *et al*, 2022; Sousa *et al*, 2023). However, how and if these kinases and transcription factors are connected within the human naïve phosphorylation network and impact biological processes remain open questions that need to be addressed by additional computational approaches. Over the past decade, numerous mechanistic modeling approaches primed by prior knowledge emerged as robust aids in comprehending the complexities of the cell signaling (Garrido-Rodriguez *et al*, 2022). These approaches use pre-existing information, annotated in public repositories (Türei *et al*, 2016; Lo Surdo *et al*, 2023; Hornbeck *et al*, 2012), about regulatory interactions among proteins, to establish a ground structure of the signaling network. Subsequently, they incorporate (multi)-omics data to generate a static representation of a specific condition (*mechanistic model*). Mechanistic models have been shown to be highly effective for studying cancer progression or drug response and for discovering novel biomarkers (Liu *et al*, 2019; Hidalgo *et al*; Massacci *et al*, 2023; Pugliese *et al*, 2023). For instance, the COSMOS pipeline has been used to generate mechanistic hypotheses from multi-omics data, including metabolomics, in patients with clear cell renal cell carcinoma (ccRCC) (Dugourd *et al*, 2021). In general, mechanistic models aim to bridge the gap between the vast omics datasets and the phenotypic outcomes observed in biological systems. However, models usually contain many nodes and edges, and this complexity hampers their functional interpretation. To tackle this issue, most of the methods use the manual exploration of the model guided by the functional enrichment analysis (Dugourd *et al*, 2021; Liu *et al*, 2019); as an alternative, other tools, such as HiPathia (Peña-Chilet *et al*, 2019), decompose pathways into functional circuits ending on phenotypes. Finally, we recently developed ProxPath (Iannuccelli *et al*, 2023), a graph-based tool designed to estimate the regulatory impact of proteins on phenotypes annotated in SIGNOR (Lo Surdo *et al*, 2023).

In this landscape, what is still missing is a strategy that integrates all these procedures (protein activity estimation, network reconstruction, and phenotypic interpretation) in a unified pipeline capable of drawing from multi-omics data a coherent picture depicting the signaling events that eventually impact hallmark phenotypes.

To fill this gap, here we present a newly implemented version (2.0) of *SignalingProfiler* (https://github.com/SaccoPerfettoLab/SignalingProfiler/), a generally applicable strategy designed to unbiasedly building mechanistic models that capture from multi-omics data signal remodeling in response to perturbations (e.g., diseases, drug treatments, etc.). *SignalingProfiler* integrates transcriptomics, proteomics, and phosphoproteomics data with the existing knowledge of molecular interactions sourced from databases such as SIGNOR (Lo Surdo *et al*, 2023) and PhosphoSitePlus (Hornbeck *et al*, 2012). The resulting model connects perturbed proteins (e.g., receptors) to effector proteins, ultimately regulating phenotypes relevant to the user’s biological context (**Figure 1**).

**Figure 1.**
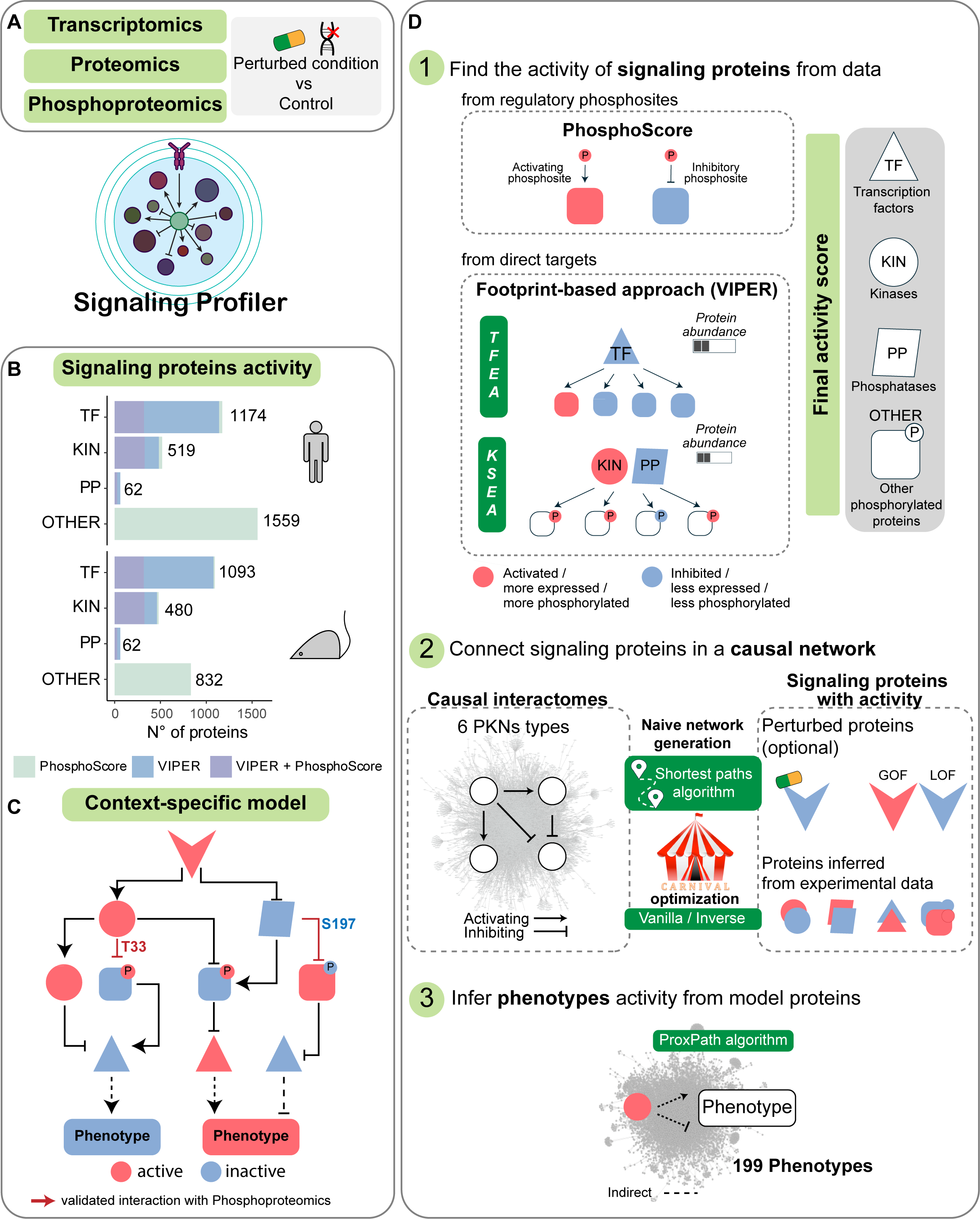
*SignalingProfiler* pipeline. **A.** *SignalingProfiler* input consists of multi-omic data collected from perturbed and control conditions (e.g., disease/ treated vs control). **B.** Coverage of *SignalingProfiler* inferable signaling proteins in human and mouse datasets, categorized by molecular function (TF transcription factors, KIN kinases, PP phosphatases, and OTHER other molecular functions). **C.** *SignalingProfiler* final output illustrates the remodeling of the signal, linking user-defined perturbed nodes (optional) with inferred proteins, and ultimately leading to relevant phenotypes. Node activities are coherent with the sign of the edges (red and blue are active and inactive proteins, respectively). Phosphoproteomics is mapped onto edges (validated interactions with phosphoproteomics). **D.** *SignalingProfiler* is a three-step modular pipeline. Step 1 derives the activity of signaling proteins from regulatory phosphosites (*PhosphoScore* method) and direct transcripts/phosphopeptides using the VIPER algorithm (*footprint-based* methods) (Alvarez *et al*, 2016). **Step 2.** A user-defined set of perturbed molecules/receptors (e.g., targets of a treatment or mutated genes in a disease) is connected to the inferred proteins using a prior knowledge network (PKN) exploiting: (i) a shortest-path algorithm to reduce the dimension of the PKN to the neighborhood of the inferred proteins (naïve network); (ii) the CARNIVAL optimization strategy (Liu *et al*, 2019) that retains only the sign-coherent interactions between proteins (context-specific network). **Step 3.** The context-specific network is connected to cellular phenotypes using the ProxPath algorithm (Iannuccelli *et al*, 2023) and the phenotype activity is obtained by integrating upstream protein activities.

The prototype of *SignalingProfiler* allowed us to uncover mechanisms of drug resistance in drug-resistant leukemia cells (Massacci *et al*, 2023; Pugliese *et al*, 2023). The current version of *SignalingProfiler* extends its utility to broader contexts, includes advanced functionalities, and incorporates expanded databases, making it a valuable resource for omics data interpretation and hypothesis generation (**Figure 1**). Here we carry out a systematic benchmarking of *SignalingProfiler* 2.0, emphasizing its advanced capabilities and broader applicability in the systems biology and network modeling fields.

## Results

### Pipeline overview

*SignalingProfiler* is an R workflow designed to unbiasedly integrate literature-derived causal networks with multi-omics data to deliver context-specific signed and oriented graphs connecting molecular entities (e.g. proteins, complexes, metabolites) and ending up on functional traits (phenotypes) (**Figure 1A-C**).

Here we provide a step-by-step description of the method.

### Step 1. Find the activity of key signaling proteins

In this step, *SignalingProfiler* derives the activity of key signaling proteins by systematically analyzing transcriptomic and (phospho)proteomic data derived from human and mouse samples. Protein activity estimation includes two main methods:

#### Footprint-based approach

Here, *SignalingProfiler* determines the activity of transcription factors, kinases, and phosphatases based on the abundance of their targets (transcripts or phosphopeptides) by integrating our newly developed algorithms and statistical tests (see Supplementary Material) with the VIPER inference method (Alvarez *et al*, 2016). This process is often referred to as Transcription Factor or Kinase Substrate Enrichment Analysis (TFEA and KSEA, respectively) (**Figure 1D, Step 1**).

The relationship between a TF/kinase/phosphatase and its specific set of transcripts/phosphopeptides is referred to as “regulon” and is extracted from public repositories (Türei *et al*, 2016; Garcia-Alonso *et al*, 2019; Lo Surdo *et al*, 2023; Müller-Dott *et al*, 2023; Johnson *et al*, 2023). A major implementation in *SignalingProfiler* 2.0 is the import of novel regulons, such as the CollecTRI resource (Müller-Dott *et al*, 2023) and the Serine Threonine Kinome Atlas (Johnson *et al*, 2023) (**Figure S1A-B**).

#### PhosphoScore

This method exploits the modulation of phosphosites in phosphoproteomics data with their impact on protein activity or stability as annotated in PhosphoSitePlus and SIGNOR (**Figure S1C-D**). Importantly, the PhosphoScore methodology allows us to extend our analysis to distinct types of molecular entities: 30% of the proteins with a regulatory phosphosite available in *SignalingProfiler* are TF/kinase/phosphatase, the remaining 70% exhibit different GO molecular functions, including, but not limited to, ubiquitin-ligase, GTP-ase, and membrane transporter activities (**Figure S1E**).

Thanks to the integration of multiple resources and the combination of PhosphoScore and footprint-based methods, the coverage of *SignalingProfiler* 2.0 is greatly expanded: a user can potentially infer nearly the entire kinome (519 and 480 kinases for human and mouse), 62 phosphatases, and over one thousand transcription factors and other signaling proteins (**Figure 1B**). Remarkably, the modular nature of the pipeline allows users to feed *SignalingProfiler* with the three datasets simultaneously (transcriptomics, proteomics, and phosphoproteomics) or with only a selection of them.

### Step 2. Connect signaling proteins in a causal network

The next step of the pipeline is the reconstruction of the molecular interactions between the modulated signaling proteins, by accessing literature-derived causal networks (**Figure 1D, Step 2**). This step includes i) the search for connections between modulated molecules detected in Step 1 within a compendium of available interactions in a prior knowledge network (PKN) and ii) the optimization of the final model.

#### The PKNs

*SignalingProfiler* 2.0 offers six categories of prior knowledge networks (PKNs), organized by organism (human or mouse) and covering signaling pathways and post-translational modifications (direct interactions) as well as gene regulation (mostly indirect interactions) derived from public resources (Lo Surdo *et al*, 2023; Hornbeck *et al*, 2012) (**Figure S2** and **Figure S3A**). Every PKN is a graph built of causal interactions represented according to the activity-flow model. Briefly, every interaction is binary, directed (has a regulator and a target of the regulation), and signed (representing either an up-or a down-regulation). The PKNs contain up to 60,807 connections (**Figure S3A**) linking a wide range of molecular entities, including proteins, fusion proteins, metabolites, and complexes (**Figure S2**).

#### The naïve network

*SignalingProfiler* allows to progressively make the PKNs context-specific, retaining only interactions in the current knowledge that are responsible for the modulation of TFs, kinases, phosphatases, and other signaling proteins (Step 1). Users have the possibility to embed in the network a set of starting perturbed nodes, which can be proteins whose activity is impacted upon genetic or pharmacological perturbation (e.g., a drug-target, a mutated protein, or ligand-stimulated receptor) **(Figure 1D, Step 2**).

First, we allow the user the possibility to remove the interactions that do not involve genes or proteins expressed in the samples (*PKN preprocessing)*. Subsequently, we provide a modular framework to identify the regulatory paths linking the perturbed nodes to transcription factors, resulting in three distinct layouts. These layouts are distinguished by the number of layers, where a layer is defined by the connection of two different molecular functions: perturbed node to kinase/phosphatase, kinase/phosphatase to other signaling protein, and other signaling protein to transcription factor (**Figure S3B**) (further details are provided in Supplementary Material).

#### The optimization

The naïve network undergoes optimization upon protein activity through the application of the Integer Linear Programming (ILP). Within the *SignalingProfiler* framework, we have incorporated two flavors of the CARNIVAL algorithm, namely Vanilla or Standard CARNIVAL (*StdCARNIVAL*) and Inverse CARNIVAL (*InvCARNIVAL*) (Liu *et al*, 2019). The CARNIVAL algorithm is developed to identify the smallest sign-coherent subnetwork, connecting as many deregulated proteins as possible. To enhance the comprehensiveness of the generated model, we have implemented a novel optimization feature that entails the execution of multiple CARNIVAL optimizations (multi-shot) for each layer of the model, producing subparts of the final model. Subsequently, these subparts are combined to form a more expansive and richer representation (**Figure S3C-F)**.

The result of these steps is a mechanistic model that can be explored at the phosphorylation-resolution level (**Figure 1C**).

### Step3. Hallmark phenotypes inference for functional interpretation

An important novelty of *SignalingProfiler* 2.0 is the implementation of the PhenoScore algorithm that infers from the model the regulation of hallmark phenotypes (**Figure 1D, Step 3)**. Specifically, it incorporates and adapts our in-house ProxPath method (Iannuccelli *et al*, 2023) (see Supplementary Material), a graph-based algorithm designed to measure the functional proximity of a list of gene products to target pathways and phenotypes, using causal interactions annotated in SIGNOR. The PhenoScore algorithm averages the activity of phenotype upstream regulators in the model and uses this value as a proxy of the activation level of phenotypes.

In summary, *SignalingProfiler* offers information on approx. 200 distinct phenotypes (e.g., Proliferation, Apoptosis, G2/M phase transition, etc.) that can be incorporated into the model (**Figure 1C**).

### Use Case

To showcase the potential of *SignalingProfiler,* we took advantage of our previously published transcriptome, proteome, and phosphoproteome dataset of breast cancer cells upon treatment with metformin (Sacco *et al*, 2016), whose molecular targets (the mammalian target of rapamycin, mTOR, and the AMP-activated protein kinase, AMPK) and phenotypic impact are well characterized (Keerthana *et al*, 2023; Garcia & Shaw, 2017; Salminen & Kaarniranta, 2012; Saxton & Sabatini, 2017; Gao *et al*, 2020; Madsen *et al*, 2015; Salani *et al*, 2014) (**Figure 2A**). Here we aim to validate whether *SignalingProfiler* can unbiasedly and systematically recapitulate from multi-omics data the metformin-induced signaling rewiring.

**Figure 2.**
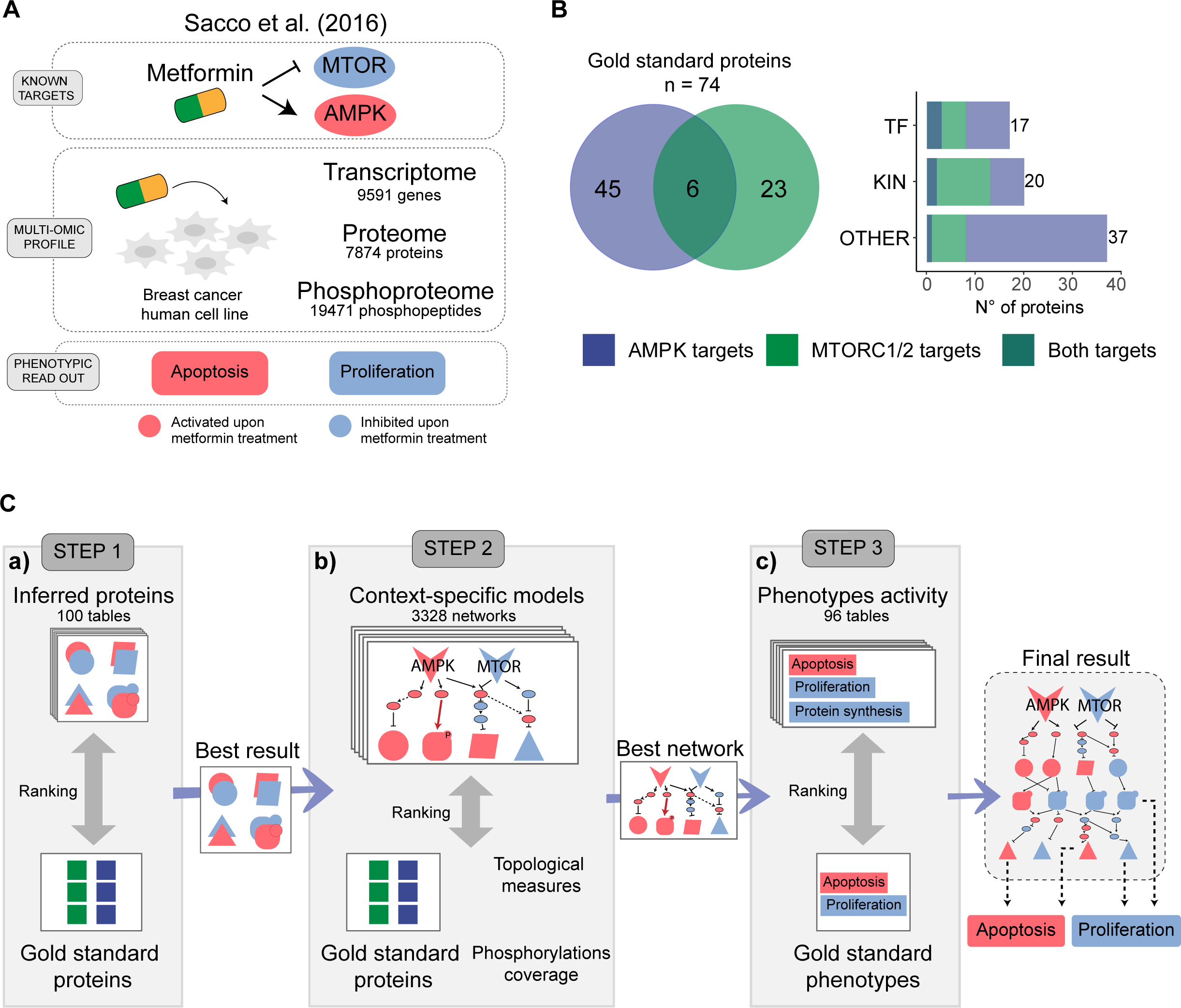
Proof-of-concept strategy. **A.** Multi-omics datasets of breast cancer cells before or after metformin treatment (Sacco *et al*, 2016). **B.** Manually curated list of 74 proteins of AMPK (blue), mTOR (green) pathways, or both (dark blue) with their expected activity after metformin treatment (*protein gold standard*). **C.** Benchmarking of the three steps of *SignalingProfiler* by testing any possible technical parameters and choosing the best result as input for the following step.

To this aim, we first manually compiled a literature-derived list of known downstream effectors and phenotypes impacted by metformin and annotated their expected activity. The so-generated *protein* and *phenotype gold standard* accounts for 74 proteins, including 17 transcription factors and 20 kinases, and 10 phenotypes (**Figure 2B, Table S1**).

This enables us to develop a standardized evaluation process, testing any possible combination of functional parameters of *SignalingProfiler* (3524 conditions), thus identifying the best parameters to set as defaults (**Figure 2C**) (see Supplementary Material).

#### Protein activity inference use case (Step 1)

Here, we inferred protein activity from experimental data of our use case (Step 1) using all possible reference databases (as sources of regulons, and of regulatory phosphosites) (**Figure S1A**) and different technical parameters.

This process yielded 100 distinct combinations of input settings and relative resulting sets of predicted protein activities (**Table S2**). Subsequently, by comparing these results with the *protein gold standard*, we systematically assessed the precision, recall, and Root Mean Squared Error (*RMSE*) associated with each combination (see Methods), to ultimately choose the most accurate and complete list of modulated proteins **(Figure 2C, panel a,** and **Table S3).**

#### Transcription factors

The procedure enabled us to infer the activity of up to 7 out of 17 (40%) transcription factors in the *protein gold standard* (**Figure 3A**). The selection of the reference database is the most impactful parameter in this task. Using combined resources like CollecTRI + SIGNOR led to a two-fold increase in the predicted TFs (**Figure 3B**). Nevertheless, when we focused on the gold standard, we observed that the regulon source maximizing both quality and coverage was Dorothea + SIGNOR (**Figure 3A**). Additionally, utilizing the hypergeometric test, which gives priority to TFs with regulons containing a high number of significantly modulated targets, tends to strengthen the signal.

**Figure 3.**
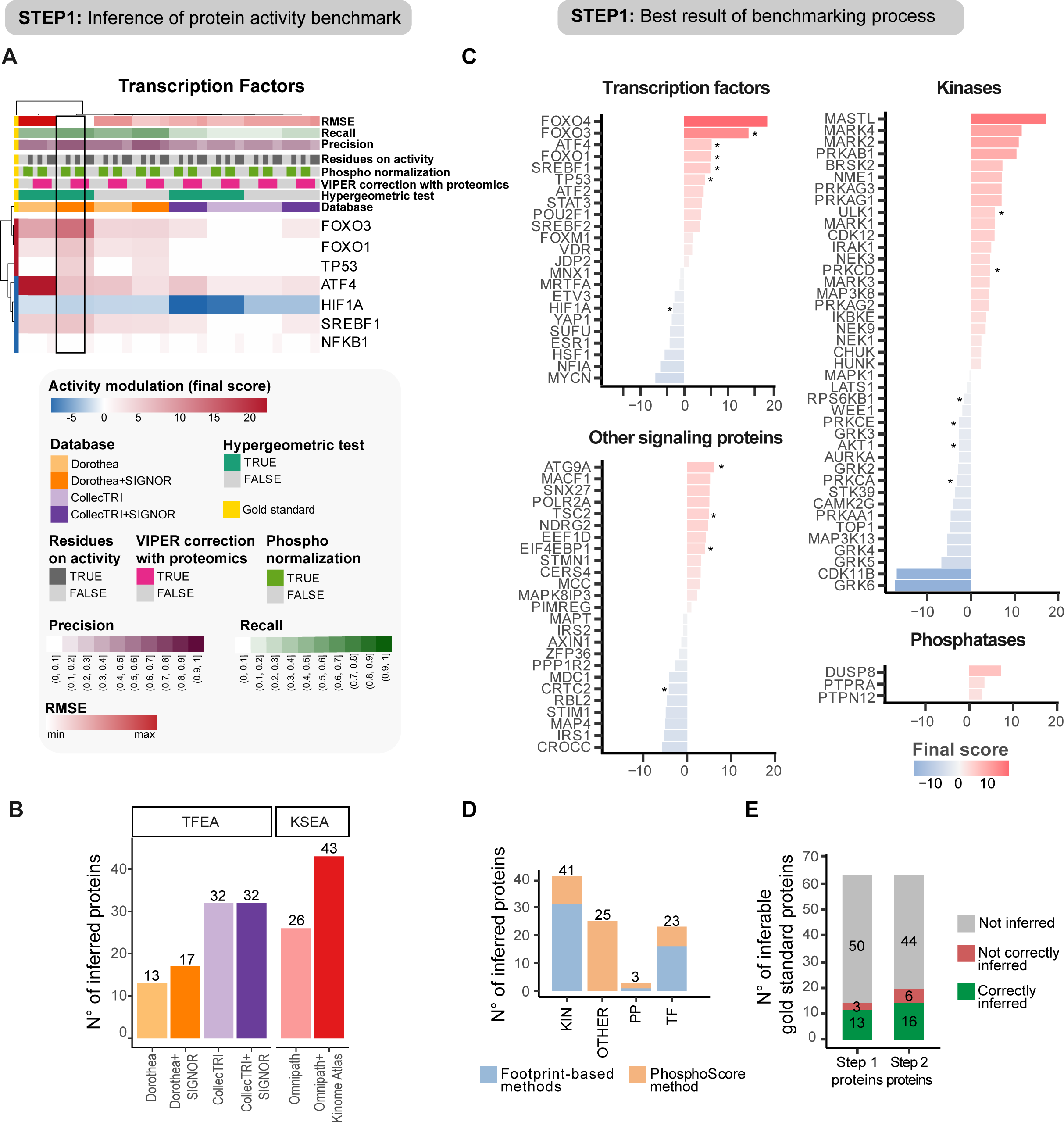
*SignalingProfiler* proteins’ activity inference benchmark and best result. A-B. Results of the benchmarking of Step 1. **A.** Heatmap reporting the inferred activity (*final score*) of gold standard transcription factors using both footprint-based and PhosphoScore methods across 64 technical conditions (see Supplementary Material). Precision, recall, and Root Mean Squared Error (RMSE) are reported for each condition by comparison with the gold standard (see Methods). The black box highlights the best technical condition set as default in *SignalingProfiler* and used in Step 2. Blue and red represent inactive and active proteins, respectively. **B.** Number of inferred proteins in KSEA or TFEA based on regulon background resources. **C.** Bar plot displaying the activity modulation (metformin-treated vs control condition) for transcription factors, kinases, phosphatases, and other signaling proteins in the top result from Step 1. Blue and red represent inactive and active proteins, respectively. **D.** Bar plot representing the proportion of transcription factors, kinases, phosphatases, and other signaling proteins identified using PhosphoScore (orange) or footprint-based methods (blue). **E.** Bar plot showing the number of proteins with inferred activity before (Step 1) and after network construction (Step 2) among the 66 inferable proteins of the gold standard.

#### Kinases

The process of inferring kinases resulted in the identification of up to 8 out of 20 (40%) kinases from the gold standard (**Figure S4A**). In general, the Omnipath and Ser/Thr Kinome Atlas integration increased the number of predicted proteins from 26 to 43, while still upholding a high level of consistency with the gold standard (**Figure 3B** and **S4A**). Moreover, the normalization of the phosphoproteomic data over proteomics (see Methods) was found to be the most influential parameter **(Figure S4A)**.

#### Other signaling proteins

Among 30 of the non-TFs/kinases in the *gold standard*, eight were identified (27%) (**Figure S4B**). This benchmarking underscores the importance of the normalization parameter and utilization of phosphosites that regulate activity rather than quantity to guarantee minimal RMSE and enhanced precision when using the PhosphoScore algorithm (**Figure S4B**).

Overall, we inferred 30-40% of the *protein gold standard*. Our inability to infer the remaining 60-70% may be attributed either to limitations in the experimental setting (e.g., drug concentration, time of treatment, choice of the breast cancer cell line) or in the pipeline itself (e.g., limited coverage in public repositories).

In summary, the best combination of parameters (**Figure S4C**) led to the inference of 23 transcription factors, 41 kinases, 3 phosphatases, and 25 other signaling proteins (**Figure 3C** and **Table S4**) and was used as an input for Step 2 **(Figure 2C, panel a)**. As expected, integrating the PhosphoScore method with footprint-based analyses expanded the number of inferred proteins **(Figure 3D**) while maintaining a high level of agreement with the gold standard (**Figure S4D**).

Remarkably, our pipeline enabled us to catch among the most highly modulated proteins many members of the gold standard (**Figure 3C**, starred proteins). In fact, among the top highly active transcription factors there is the Forkhead box O family (FOXO family) which is directly affected by the metformin-activation of AMPK complex (Queiroz *et al*, 2014; Greer *et al*, 2009). Coherently, Hypoxia Inducible Factor 1 Subunit Alpha (HIF1A), an indicator of hypoxia, was among the top down-regulated TFs (Zhou *et al*) along with N-myc proto-oncogene (MYCN), a marker for cell proliferation (Wang *et al*, 2014) (**Figure 3C**, transcription factors panel). Furthermore, the PhosphoScore algorithm allowed us to observe the inhibition of Insulin Receptor Substrate 1 and 2 (IRS1 and IRS2), a phenomenon already associated with metformin-induced AMPK activation (Zakikhani *et al*, 2010) (**Figure 3C**, other signaling proteins panel).

#### Network construction use case (Step 2)

A key challenge in multi-omics data integration is extracting the cause-effect relationships underlying the experimental data. Translated to our use case, this task attempts to address the specific molecular events triggered by metformin treatment. To this aim, we extracted the direct and indirect connections linking the mTOR protein and AMPK complex to the proteins modulated in their activities through any possible framework in Step 2 of *SignalingProfiler* (**Figure S3**). This process involved the screening and the evaluation of 3328 possible resulting networks, by ranking them according to a combined score, that considers elements such as the consistency with the protein gold standard and topological graph metrics (see Methods) **(Figure 2C, panel b, S5, S6**, and **Table S5).**

Overall, the average computation time of the analyses was 200 seconds (**Figure S5A**). The networks were obtained from 2989 runs over 3328, with most models consisting of only one component (**Figure S5B**) and demonstrating a strong fit to the power law (**Figure S5C**). As shown, the integration of the Ser/Thr Kinome Atlas into the prior knowledge network (see Methods), as well as the usage of two- and three-layer naïve network types, led to an increased computation time and dimensionality and, as expected, an increased coverage of metformin-dependent phosphorylation events (**Figure S6A-L**). Unexpectedly, we found out that no major differences were detected between the two- and the three-layered networks, as revealed by the evaluation of the path length from mTOR and AMPK to endpoints in the signaling cascade (**Figure S6N, Q**).

We also benchmarked the two types of CARNIVAL differentiated by the usage of starting perturbed nodes as constraints. The *invCARNIVAL* requires increased computational time (**Figure S6C)** and returns smaller networks (**Figure S6F, I)**. Moreover, due to the limited constraints and the complexity of the basic network, only 3% of the models generated by *inv*CARNIVAL correctly inferred both mTOR and AMPK, whereas 55% of them inferred only one of them.

On the other hand, the *stdCARNIVAL* returns larger networks with the two-shot optimization outperforming the one- and three-shot ones, in the number of nodes and phosphorylation events **(Figure S6F, I, L)**, without increasing the computation time **(Figure S6C)**.

Overall, the quality of the models with respect to the gold standard was satisfactory, with an average precision (or specificity) and recall (or sensitivity) of 0.75 and 0.35, respectively (**Figure S5D, E)**.

We ranked the models according to the combined score (**Figure S5F** and **Table S5**) and set as default the most frequent values of parameters in the top 100 models (**Figure S7A)**. The top-quality network (Network1554) accounts for 99 nodes and 219 edges, which include new proteins of the gold standard (**Figure 3E)** and recapitulates the expected mTOR pathway inactivation and AMPK pathway activation upon metformin treatment (**Figure 4A**). The gold standard proteins form a highly interconnected submodule within the final network (**Figure S7B**), however accounting for only 22% of its nodes. This indicates that the model not only recapitulates known mechanisms but also proposes new ones, including the modulation of MAPK and CDK pathways (**Figure 4A**).

**Figure 4.**
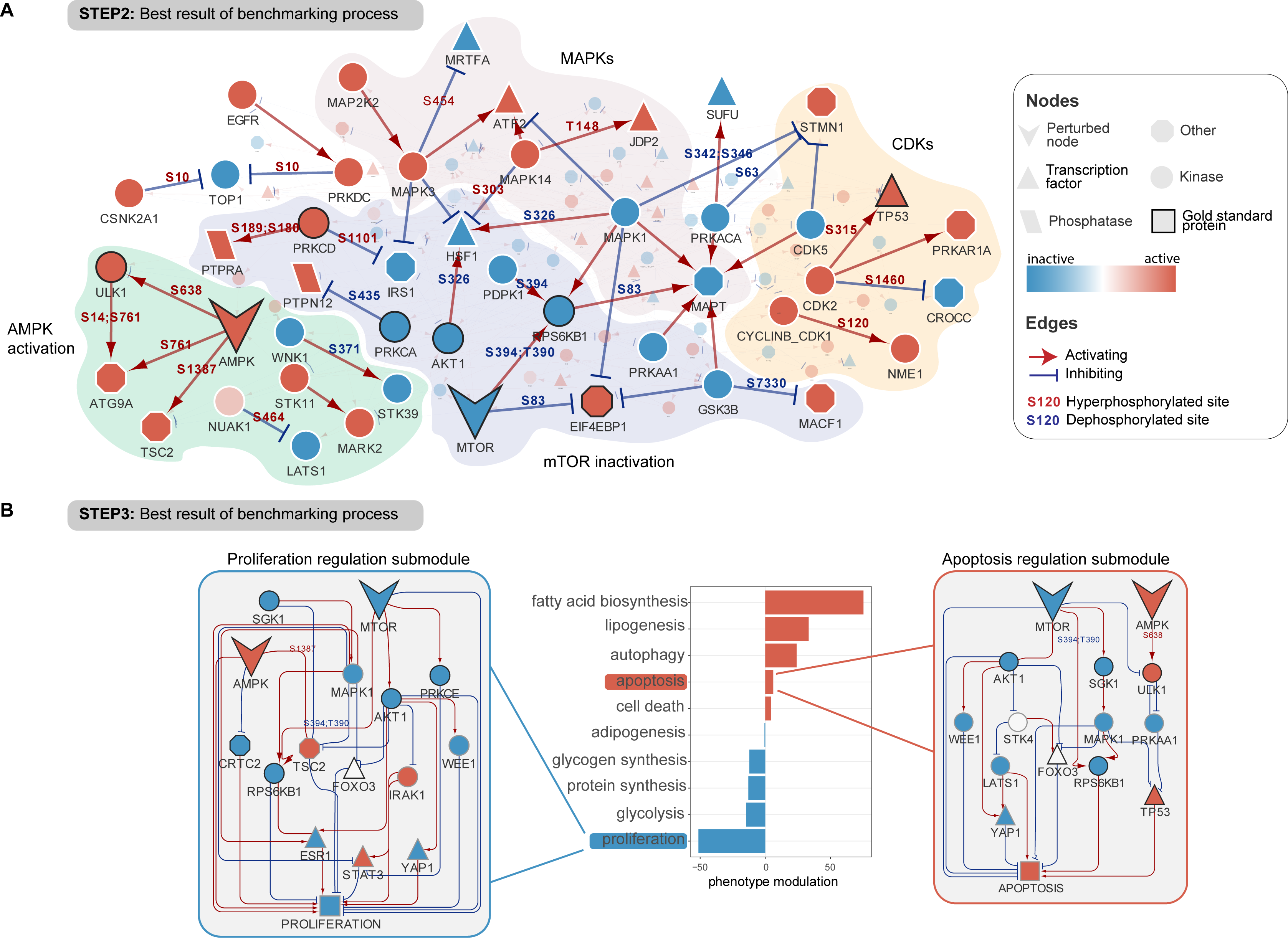
Optimal *SignalingProfiler* result of metformin-induced signaling rewiring reconstruction and its phenotypic impact. **A.** Visualization of the best model (**Figure S9**) of the benchmarking of Step 2 focused on proteins involved in phosphorylation events significantly deregulated between metformin and control. Nodes and edges are displayed according to the legend. Blue and red nodes represent inactive and active proteins, respectively. Background areas highlight subnetworks associated with known signaling pathways. **B.** Bar plot reporting the inferred modulation of phenotypes upon metformin treatment after Step 3 of *SignalingProfiler* (activation in red, inhibition in blue), and the functional circuits extracted from the final model connecting AMPK and mTOR to apoptosis and proliferation.

#### Phenotype inference use case (Step 3)

Finally, we used the top-quality network derived from Step 2 as input for the inference of phenotypic outcomes. To note, the PhenoScore algorithm considers various modalities, resulting in a total of 96 potential outcomes (**Figure 2C, panel c, S8A,** and **Table S6**). We developed an aggregated score to quantify the accuracy of prediction with respect to the *phenotypic gold standard* and ranked the results accordingly. The top 10 parameters were set as default for the PhenoScore algorithm (**Figure S8B).** Then, we selected the most accurate prediction of the *phenotypic gold standard* (**Figure S8C**), and we created a final model (109 nodes and 298 edges, **Figure S9**, and **Table S7**) depicting the metformin-induced signaling axes impacting the selected phenotypes (**Figure 4B**). Interestingly, in this final model, metformin results in the activation of death-associated pathways (e.g., apoptosis and cell death) and autophagy (the most characterized phenotypic hallmark of mTOR inhibition (Gao *et al*, 2020) (**Figure S8D**), and in the inhibition of proliferation and biosynthetic pathways (e.g., protein synthesis) (**Figure 4B and S9)**.

In summary, the findings from this study demonstrate that *SignalingProfiler* is a powerful tool for extracting molecular hypotheses from multi-omics data upon a perturbation (**Figure 4A**) and identifying functional circuits that impact phenotypes (**Figure 4B**).

## Discussion

In this paper, we thoroughly present *SignalingProfiler* 2.0, a method to systematically create mechanistic context-specific networks of signaling remodeling and to identify functional circuits impacting phenotypes.

Here, we show that *SignalingProfiler* 2.0 is a modular pipeline that allows users (i) to unbiasedly derive the activity of proteins from the integration of proteogenomic data with prior knowledge information deposited in public repositories; (ii) to connect the identified proteins to generate a coherent network that explains nodes’ change in activity; (iii) to estimate the activation level of hallmark phenotypes and integrate them in the final model. With respect to the first prototype (Massacci *et al*, 2023), *SignalingProfiler* 2.0 now incorporates extended background databases, such as CollectTRI (Müller-Dott *et al*, 2023) and Ser/Thr Kinome Atlas (Johnson *et al*, 2023), increasing the coverage and the accuracy of protein activities’ prediction. Indeed, it is possible to estimate the activity of more than 3300 proteins, including, but not limited to, kinases, phosphatases, and transcription factors. Novel implementations of *SignalingProfiler* 2.0 include PKN browsing methods, optimization strategies, and PhenoScore inference with ProxPath (Iannuccelli *et al*, 2023). In particular, the PhenoScore makes it possible to map up to 200 cellular phenotypes onto the final model. This represents an effective strategy of feature reduction and a valuable resource for omics data interpretation and hypothesis generation in diverse biological contexts.

To systematically benchmark the ability of *SignalingProfiler* 2.0 to recapitulate the effect of a well-characterized drug on signaling rewiring, we took advantage of a multi-omics dataset derived from metformin-treated breast cancer cells. The result is a hierarchical mechanistic network, accounting for 109 nodes and 309 edges, and incorporating mTOR and AMPK known down-stream effectors (e.g., FOXOs, PRKs) as well as novel promising signaling axes (e.g., AKT1-WEE1 axis) by which metformin can exert its anti-cancer activity. In this analysis, we assess the performance and the robustness of our approach, and we set the default parameters by prioritizing quality rather than quantity. Importantly, we observe that different parameter selections might change the coverage and the strength of the solutions predicted by the approach, but this never leads to an opposite estimated activity (**Figure 4A** and **S4**), thereby demonstrating the overall robustness of the method.

An important feature of *SignalingProfiler* 2.0 is its flexibility. Flexibility on available multi-omics data (e.g. users can employ only transcriptomic or phosphoproteomic data); flexibility on the organism choice (mouse and human data are accepted) and flexibility on the type of perturbed nodes, since we include relations that are both signaling or transcriptional. Importantly, thanks to its modular structure, users have the possibility to use only a limited number of steps of the pipeline and, possibly, to integrate *SignalingProfiler* 2.0 with other methods for protein activity estimation and network optimization.

Indeed, *SignalingProfiler* 2.0 is not the sole method to generate a mechanistic network from omics data. Here, we report a systematic comparison of *SignalingProfiler* 2.0 with a panel of similar methods, released from 2017 to 2022 (Dugourd *et al*, 2021; Liu *et al*, 2019; Köksal *et al*, 2018; Babur *et al*, 2021; Bradley & Barrett, 2017; Browaeys *et al*, 2020; Fortelny & Bock, 2020; Peña-Chilet *et al*, 2019) (**Table 1**). Our analysis reveals that *SignalingProfiler* 2.0 is: (i) one of the few techniques directly annotating meta-information about the molecular function at node/protein levels, (ii) is the sole tool capable of estimating the activity of proteins, aside from kinases and phosphatases, from the phosphoproteomic data and (iii) is the sole approach together with CausalPath (Babur *et al*, 2021)combining proteomics in the analysis and, apart from HiPathia (Peña-Chilet *et al*, 2019), integrating phenotypes with their activation status into the ultimate model to unbiasedly derive functional circuits.

**Table 1.** Qualitative comparison of *SignalingProfiler* and existing methods.

|  |  | SignalingProfiler 2.0 | COSMOS | CausalPath | TPS | CARNIVAL | HiPathia | CausalR | NicheNet | KPNN |
| --- | --- | --- | --- | --- | --- | --- | --- | --- | --- | --- |
| **Omics data properties** |  |  |  |  |  |  |  |  |  |  |
| **Omics layers** | Transcriptomics | X | X | X |  | X | X | X | X | X |
|  | Proteomics | X |  | X |  |  |  |  |  |  |
|  | Phosphoproteomics | X | X | X | X |  |  |  |  |  |
|  | Metabolomics |  | X |  |  |  |  |  |  |  |
| **Biological resolution** | Bulk | X | X | X | X | X | X | X | X | X |
|  | Single-cell |  |  |  |  |  | X |  | X |  |
| **Omics data used as** | Measurements (observations) |  |  |  | X |  | X |  |  | X |
|  | Statistical scores (contrast/correlation) | X | X | X | X | X |  | X | X |  |
| **Additional input** | Does not use additional inputs |  | X |  |  |  | X | X |  | X |
|  | Accepts additional inputs | X |  | X |  | X |  |  | X |  |
|  | Requires additional inputs |  |  |  | X |  |  |  |  |  |
| **PKN properties** |  |  |  |  |  |  |  |  |  |  |
| **Sign** | Directed |  |  |  | X |  |  |  | X | X |
|  | Activations/Inhibitions | X | X | X | X | X | X | X |  |  |
| **Size** | Pathways |  |  |  |  |  | X |  |  |  |
|  | Large networks | X | X | X | X | X |  | X | X | X |
| **Biological content** | Protein signaling interactions | X | X | X | X | X |  |  | X | X |
|  | Gene regulatory interactions | X | X | X |  |  | X |  | X | X |
| **Method properties** |  |  |  |  |  |  |  |  |  |  |
| **Omics to PKN** | Direct mapping to nodes |  |  | X | X |  | X | X | X | X |
|  | Indirect mapping to nodes | X | X |  |  | X |  |  |  |  |
| **Not-measured nodes** | Estimates unmeasured nodes state | X | X |  | X | X | X | X |  | X |
|  | Includes unmeasured nodes in output | X | X |  | X | X | X | X | X | X |
| **Algorithm type** | Edge filtering and shortest path | X |  | X |  |  |  | X | X |  |
|  | Recursive signal propagation and heat diffusion |  |  |  |  |  | X |  |  |  |
|  | Integer linear programming | X | X |  | X | X |  |  |  |  |
|  | Neural networks |  |  |  |  |  |  |  |  | X |
| **Final network properties** |  |  |  |  |  |  |  |  |  |  |
|  | Includes user-defined perturbed nodes | X |  |  |  | X | X |  |  |  |
|  | Includes phenotypes | X |  |  |  |  | X |  |  |  |
|  | Includes meta-information on nodes | X | X |  |  | X |  |  | X |  |
|  | Includes meta-information on edges regarding phosphorylation events | X |  | X | X |  |  |  |  |  |

Finally, the comparison with other methods highlights some of the limitations of our pipeline. As compared to tools such as COSMOS (Dugourd *et al*, 2021), *SignalingProfiler* 2.0 does not include metabolomic data. At the present state, additional types of regulation such as epigenetic acetylomic and ubiquitylomic data, which are becoming more popular (Li *et al*, 2022; Aslanyan *et al*, 2023) cannot be integrated into the signaling and represent a future challenge to face. Also, as for all the methods that base their prediction on prior knowledge, *SignalingProfiler* 2.0 suffers from the limited coverage of available information in public repositories: either regulon databases and causal interaction resources are incomplete and offer information for less than 50% (about 9,000 proteins) of the Uniprot-SwissProt proteome.

In summary, *SignalingProfiler* 2.0 is a versatile and flexible pipeline that efficiently generates mechanistic networks from multi-omics data hierarchically bridging signaling molecules to phenotypic traits. As such, it addresses the emergent need to extract interpretable networks and derive biologically relevant information from complex multi-omics data. We expect that in the multi-omics era, where the proteogenomic characterization of human samples and biopsies are becoming increasingly more available to the public (Ng *et al*, 2022; Rudnick *et al*, 2016), *SignalingProfiler* 2.0 could pave the way to the development of personalized medicine strategies.

**Recommendation for use:** *SignalingProfiler* can be applied to the results of any experiments having at least transcriptomics or phosphoproteomics to identify signaling rewiring that is supported by literature knowledge. To use *SignalingProfiler*, the user has to provide a fold-change between the conditions under investigation, and measurement values need to be associated with related primary gene symbols and UniProt ID. Moreover, the phosphopeptide measurements need to specify the phosphorylation site with respect to their canonical UniProt sequence, in a specific format described in the GitHub page. The user can choose to visualize the whole network, the network focused on the significant phosphorylation events (*phospho-layout)* or given one or more phenotypes the impacting functional circuits (*pheno-layout*).

## Acknowledgments

We thank Prof. Gianni Cesareni and Prof. Luisa Castagnoli for their fruitful discussion during manuscript revision. Dr. Serena Paoluzi and Dr. Marta Iannuccelli for their technical support. This research was funded by the Italian Association for Cancer Research (AIRC) with a grant to L.P. (MFAG Grant n. 28858) and a grant to F.S. (Start-Up Grant n. 21815). Also, L.P. and F.S. are supported by a joint PRIN 2022 PNRR grant (n. P2022JRETW), funded by the European Union - NextGenerationEU. V.V. is supported by PON-MUR fellowship (n. DOT13IEP1U-1)

## Author contributions

Conceptualization, V.V., L.P., F.S.; methodology, V.V., L.P., F.S.; formal analysis, V.V; investigation, V.V.; writing original draft preparation, V.V., F.S., L.P.; resources, F.S., L.P.; writing review and editing, all; supervision, F.S., L.P.; funding acquisition, F.S., L.P. All authors have read and agreed to the published version of the manuscript.

## Material and Methods

### PKNs creation

We downloaded all causal interactions available for *Mus musculus* (TaxID = 10090) and *Homo sapiens* (TaxID = 9606) from the SIGNOR and PhosphoSitePlus® resources. SIGNOR 3.0 datasets, retrieved via the REST API, are based on information up to November 2023. Interactions labeled ‘down-regulates,’ ‘up-regulates,’ and ‘form complex’ in SIGNOR were assigned values of −1, 1, and 1, respectively. Interactions involving entities with the TYPE ‘protein family’ in SIGNOR were excluded. Causal phosphorylations from PhosphoSitePlus® were obtained by manually downloading and combining two independent tables: kinase-phosphosite interactions (’Kinase_Substrate_Dataset.gz’) and the regulatory role of phosphosites on proteins (’Regulatory_sites.gz’). The tables were joined using the UniProt ID and modified residue as keys. The content of the ‘ON_FUNCTION’ column in PhosphoSitePlus® representing the regulatory role of phosphosites was mapped to values of 1, −1, or 0. These manipulated datasets were merged and filtered to retain interactions with a defined regulatory effect (−1 or 1). For *Homo sapiens*, causal interactions derived from the Ser/Thr Kinome Atlas were added to SIGNOR and PhosphositePlus datasets (see Methods ‘Ser/Thr Kinome Atlas parsing’). UniProt IDs were updated, and primary Gene Names were retrieved using the UniProt database’s REST API. The primary Gene Names of the involved entities were used as keys for each interaction, and multiple UniProt IDs and attributes (e.g., TYPE, DATABASE field of SIGNOR) were collapsed into a single string. We created six Prior Knowledge Networks (PKNs) adding increasingly exclusive filtering criteria: no filtering (PKN2 for human and PKN6 for mouse), removal of indirect interactions representing ‘transcriptional regulations’ (PKN1 for human and PKN5 for mouse), removal of Kinome Atlas interactions (PKN4), and removal of direct interactions not involving proteins (PKN3). The number of nodes and edges of each PKN is shown in **Figure S2** and **S3**.

### Ser/Thr Kinome Atlas parsing

We obtained Supplementary Table 3 from the work of (Johnson *et al*, 2023), containing information on 89752 serine (Ser) and threonine (Thr) sites and their probabilities (or percentile) of being phosphorylated by 303 Ser/Thr kinases. We kept phosphosite-kinase relations with a percentile higher than 88 (“*regulon threshold”*) and 99 (“*PKN threshold”),* retaining 3,134,109 and 291,682 relations, respectively.

The “*regulon threshold”* of 88 was determined as the median value from the distribution of percentiles of phosphosite-kinase relations documented in SIGNOR or PhosphoSitePlus®. These relations were incorporated into the regulons for kinase inference analysis, with weights assigned proportionally to the percentiles within the range of 0.5 to 0.9.

The “*PKN threshold”* of 99 was chosen to keep only the most accurate relationships. We joined this table with the PhosphoSitePlus table on the regulatory role of phosphosites (’Regulatory_sites.gz’) using the phosphosite as key. As a result, we included 28,012 interactions in the prior knowledge networks, representing relationships between kinases from the Atlas and proteins for which the regulatory phosphosite is known.

Each kinase was annotated with its UniProt ID.

### Benchmarking strategy

#### Metformin multi-omics dataset preparation

We downloaded relevant tables from our work as published in (Sacco *et al*, 2016) to build transcriptomic, proteomic, and phosphoproteomic data tables. The so-obtained information was parsed and adapted to make it *SignalingProfiler* compliant.

Briefly, the dataset accounted for 9591, 7974, and 15812 quantified transcripts, proteins, and phosphosites. These tables included computed fold-change values among three replicates of both the control and metformin conditions.

#### Normalization of phosphoproteomics over proteomic data

We created a normalized phosphoproteomic dataset by adjusting the fold-change in phosphorylation in response to metformin treatment based on the corresponding fold-change in protein abundance. To achieve this, we calculated the difference between the phosphorylation level of the phosphosite and its associated fold-change in protein abundance.

The phosphorylation levels of phosphosites that showed modulation in proteomics with the same direction were reduced, while those with opposite phosphorylation and protein abundance changes were increased. We then computed the Z-score for the new distribution of phosphorylation fold-changes using their mean and we defined corrected fold-changes with an absolute value higher than 1.96 (i.e., p-value < 0.05) significant.

#### Protein and phenotypic gold standard creation

A list of 74 proteins with their expected activity modulation to metformin treatment (*protein gold standard*) was compiled from three recent papers (Keerthana *et al*, 2023; Garcia & Shaw, 2017; Salminen & Kaarniranta, 2012; Saxton & Sabatini, 2017) focusing on mTOR and AMPK pathways. Since metformin inhibits mTOR (and activates AMPK), negative and positive targets of mTOR (and AMPK) were set to active and inactive (inactive and active), respectively. Each protein was manually cross-referenced and converted to its primary gene name. The molecular function of each protein was annotated using *SignalingProfiler*. The resulting gold standard protein list was compared with proteins in *SignalingProfiler* databases, including TFEA or KSEA regulons and the PhosphoScore database. Notably, eight proteins were not found in the databases and were consequently labeled as ‘not inferable’ proteins. Additionally, a list of 10 phenotypic traits with their expected modulations upon metformin treatment (*phenotypic gold standard*) was compiled, based on the phenotypic readout from our previous work (Sacco *et al*, 2016) and three referenced papers (Gao *et al*, 2020; Madsen *et al*, 2015; Salani *et al*, 2014).

The complete gold standard dataset is available in **Supplementary Table 1**.

#### SignalingProfiler benchmarking

We ran the *SignalingProfiler* pipeline with all technical parameter combinations. A detailed explanation of *SignalingProfiler* functions and parameters is provided in Supplementary materials. Briefly, any test combination of parameters was evaluated by measuring precision, recall, and Root Mean Squared Error (RMSE) using protein and phenotypic gold standard lists.

#### Precision, recall, and RMSE definition

We defined an inferred protein matching and diverging the expected value, as *true* and *false positive*, respectively*. False negatives* were proteins present in the gold standard but not inferred. We calculated as quality metrics (i) *precision*, the ratio of true positives to the sum of true and false positives, (ii) *recall,* the ratio of true positives to the sum of true positives and false negatives, and (iii) *RMSE,* the squared mean difference between predicted and expected values. The eight ‘not inferable’ gold standard proteins were not considered in the quality metrics computation.

#### Step 1 benchmarking

*SignalingProfiler* independently infers the activity of transcription factors, kinases/phosphatases, and phosphorylated proteins. Transcription factors/kinases/phosphatases can be inferred with footprint-based methods, PhosphoScore, or a combination of both. The parameter combinations for the inference of transcription factors, kinases/phosphatases, and phosphorylated yielded 64, 32, and 4 results, respectively (**Supplementary Table 2**). Each result referred to a unique selection of parameters for the type of regulons/phosphosites database, the usage of the Hypergeometric Test, VIPER correction with proteomics; and correction of phosphoproteomics over proteomics.

The default setting for Step 1 was determined by selecting the result for each molecular function that maximizes precision and recall while minimizing RMSE (**Supplementary Table 3 and 4**).

#### Step 2 benchmarking

The network construction step involves 10 parameters (see Supplementary Material for details). All combinations resulted in 3328 possible results, but only 2989 combinations yielded actual networks. Each model was annotated with *computation* time (sum of naïve network computation and CARNIVAL optimization time); *topological* metrics (nodes, edges and components, clustering coefficient, diameter, fit to the power law, maximum path length between end nodes and AMPK, mTOR and Perturbation node created by Inverse CARNIVAL); *biological* metrics, such as the precision, recall and RMSE with respect to the gold standard, and the number of interactions validated by quantified or significant experimental phosphorylations (**Supplementary Table 5**). We developed a **Step 2 combined score** defined as follows:

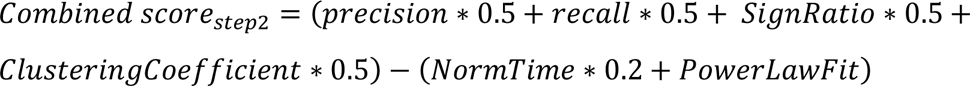

where *SignRatio* is the ratio between the number of edges that are validated by significant phosphorylation events over the total and the *NormTime* is the ratio between each computation time and its maximum. The best model was selected based on the highest aggregate score (Network1554 with 99 nodes and 219 edges). We set the default parameters for Step 2 by choosing the most frequent parameters’ values among the top 100 models.

#### Step 3 benchmarking

The benchmarking of phenotypic traits inference considered 6 different parameters (see Supplementary Material for details). The ten phenotypes of the phenotypic gold standard were selected, including apoptosis, autophagy, adipogenesis, biosynthesis of fatty acids, glycogen and proteins, proliferation, and glycolysis. We obtained 96 results that were compared to the phenotypic gold standard, and we annotated precision, recall, RMSE, and computation time (**Supplementary Table 6**). We formulated a **Step 3 combined score**:

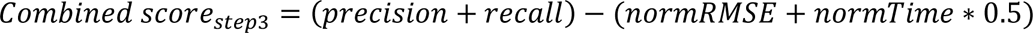

where *normRMSE* and *normTime* are the ratio of its value and its maximum.

We linked the phenotypes’ values with the highest combined score to their regulators in the Step 2 model, resulting in a final optimized network of 109 nodes and 309 edges (**Supplementary Table 7**).

The network is publicly available for browsing at: https://www.ndexbio.org/viewer/networks/fa22e724-b54b-11ee-8a13-005056ae23aa. The Step 3 default was set by considering the most represented parameters’ values among the top 10 results.

### *SignalingProfiler* output visualization

The optimized network generated by *SignalingProfiler* was displayed on Cytoscape using the RCy3 package (v. 2.14.2). Two XML files provided within the *SignalingProfiler* R package were used to visualize the network in Cytoscape.

The *“SignalingProfiler layout”* provides users with a clear and intuitive visual representation of the entire model (used in **Figure 4B, S7B, and S9**). On the other hand, the *“Phosphorylation layout”* (used in **Figure 4A**) allows users to focus specifically on proteins involved in experimentally confirmed phosphorylation events.

## Code availability

All code used for *SignalingProfiler* benchmarking is available at https://github.com/SaccoPerfettoLab/SignalingProfiler_Benchmarking/. *SignalingProfiler* 2.0 R package code is available at https://github.com/SaccoPerfettoLab/SignalingProfiler.

