## Supplementary Material and Methods for "*SignalingProfiler* 2.0: a network-based approach to bridge multi-omics data to phenotypic hallmarks"

### Supplementary material for Venafra et al.

#### Supplementary Figures

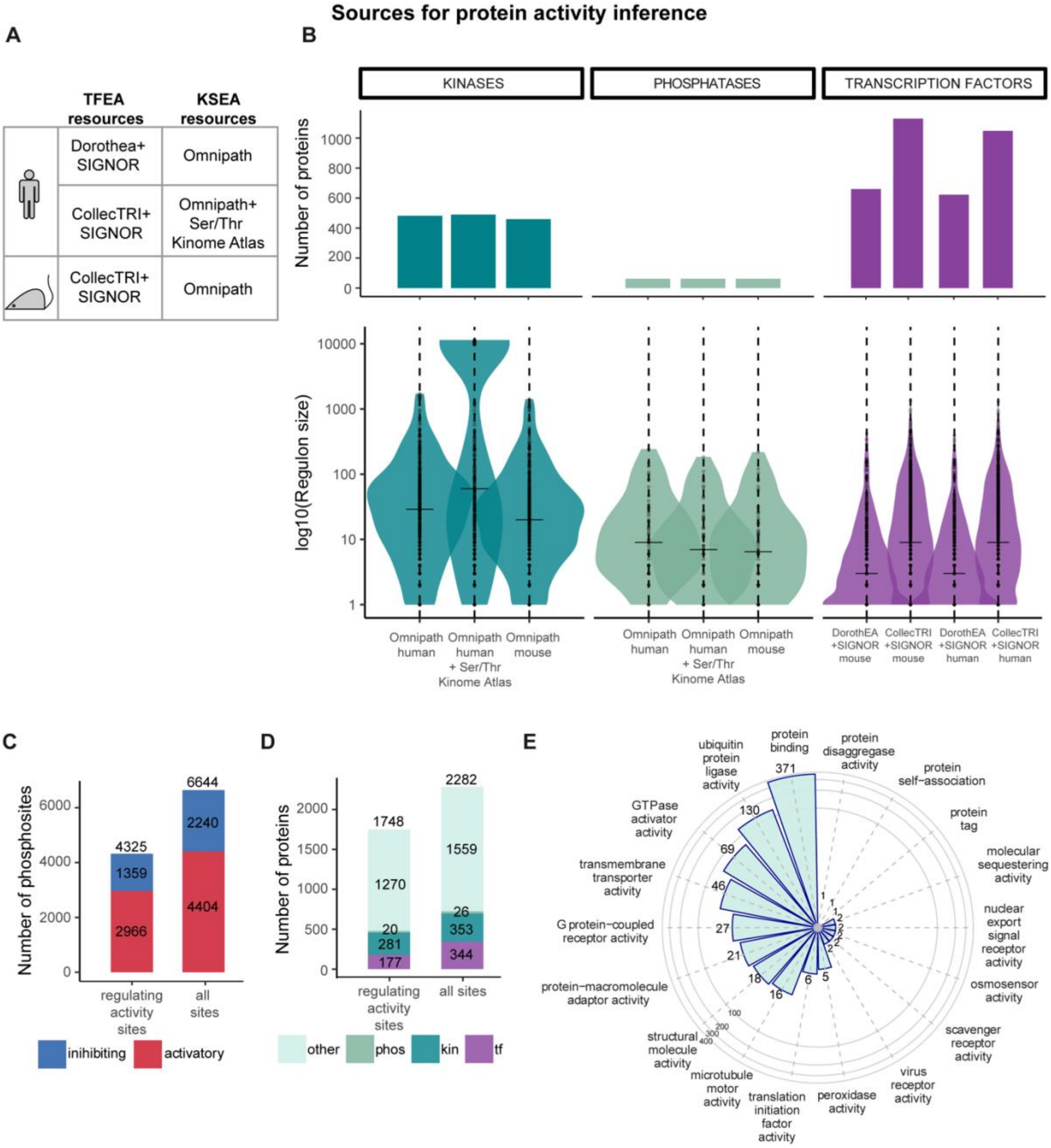


**Figure S1.** ***SignalingProfiler* sources for protein activity inference.**

**A-B.** *SignalingProfiler* regulon sources for footprint-based activity inference of transcription factors (transcription factor enrichment analysis or TFEA) and of kinases and phosphatases (kinase substrates enrichment analysis or KSEA) (**A**) and their associated number of inferable proteins (upper panel) and distribution of the regulon size (lower panel) (**B**).

**C-D.** Number of *SignalingProfiler* phosphosites regulating protein activity or stability (**C**) and their associated proteins (**D**), as obtained from SIGNOR (Lo Surdo *et al*, 2023) and PhosphoSitePlus (Hornbeck *et al*, 2012), for the PhosphoScore analysis.

**E.** GO molecular functions, obtained with gProfiler (Goel *et al*, 2012), associated with proteins having regulatory phosphosites in the PhosphoScore database and that are not kinases, phosphatases, or transcription factors and defined in *SignalingProfiler* as "other signaling proteins" (or OTHER).

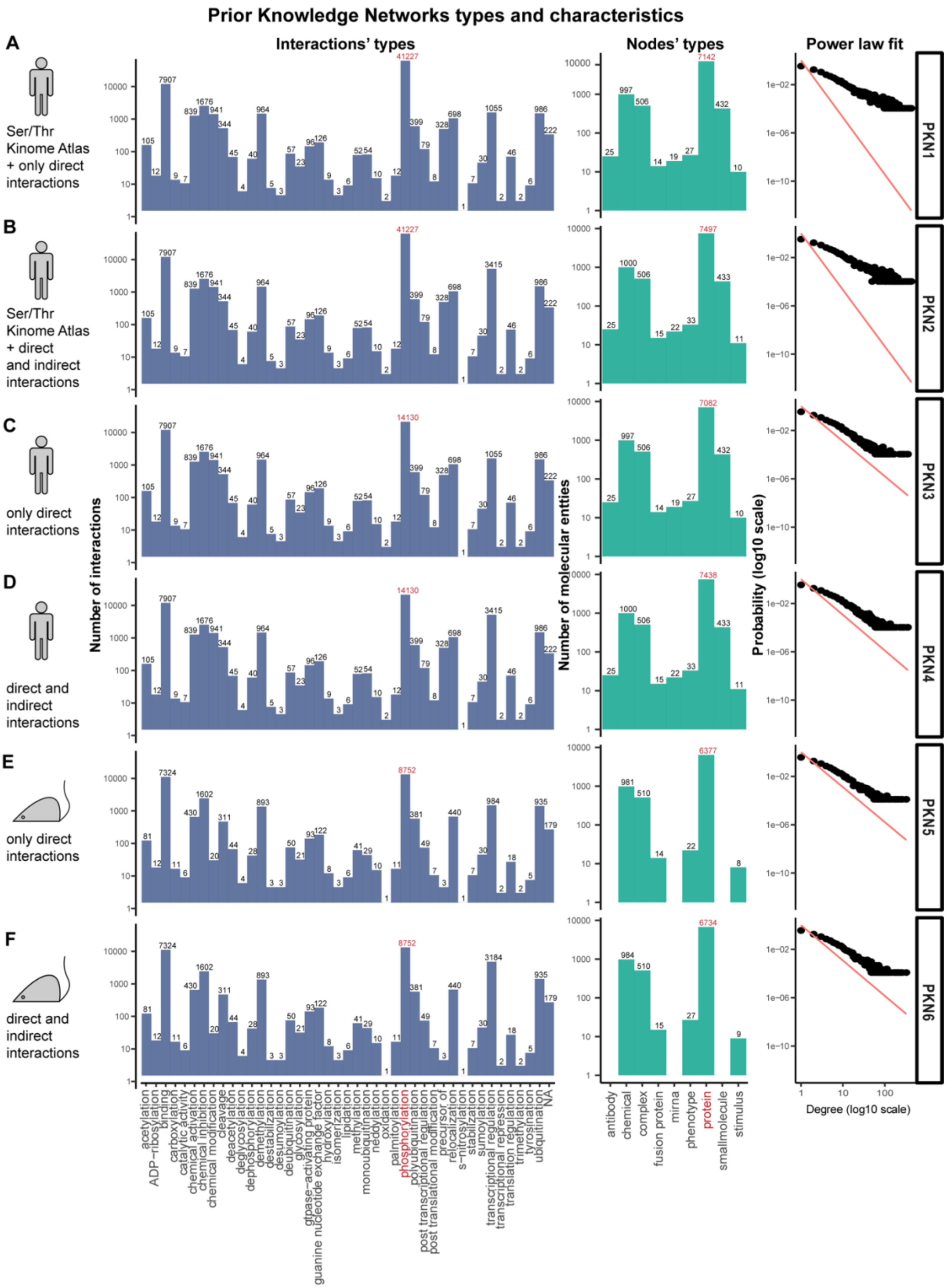


**Figure S2. The six Prior Knowledge Networks (PKNs) of *SignalingProfiler*.**

**A-F.** For each PKN: description reporting organism, integrated resource, and type of interactions (*left panel*); bar plot representing the number of interactions for each type of molecular event (*mid-left panel*); bar plot representing the number of molecular entities for each type (*mid-right panel*); scatterplot of the distribution of the PKN nodes degree in log10 scale and its fit to the power law (red line) (*right panel*).

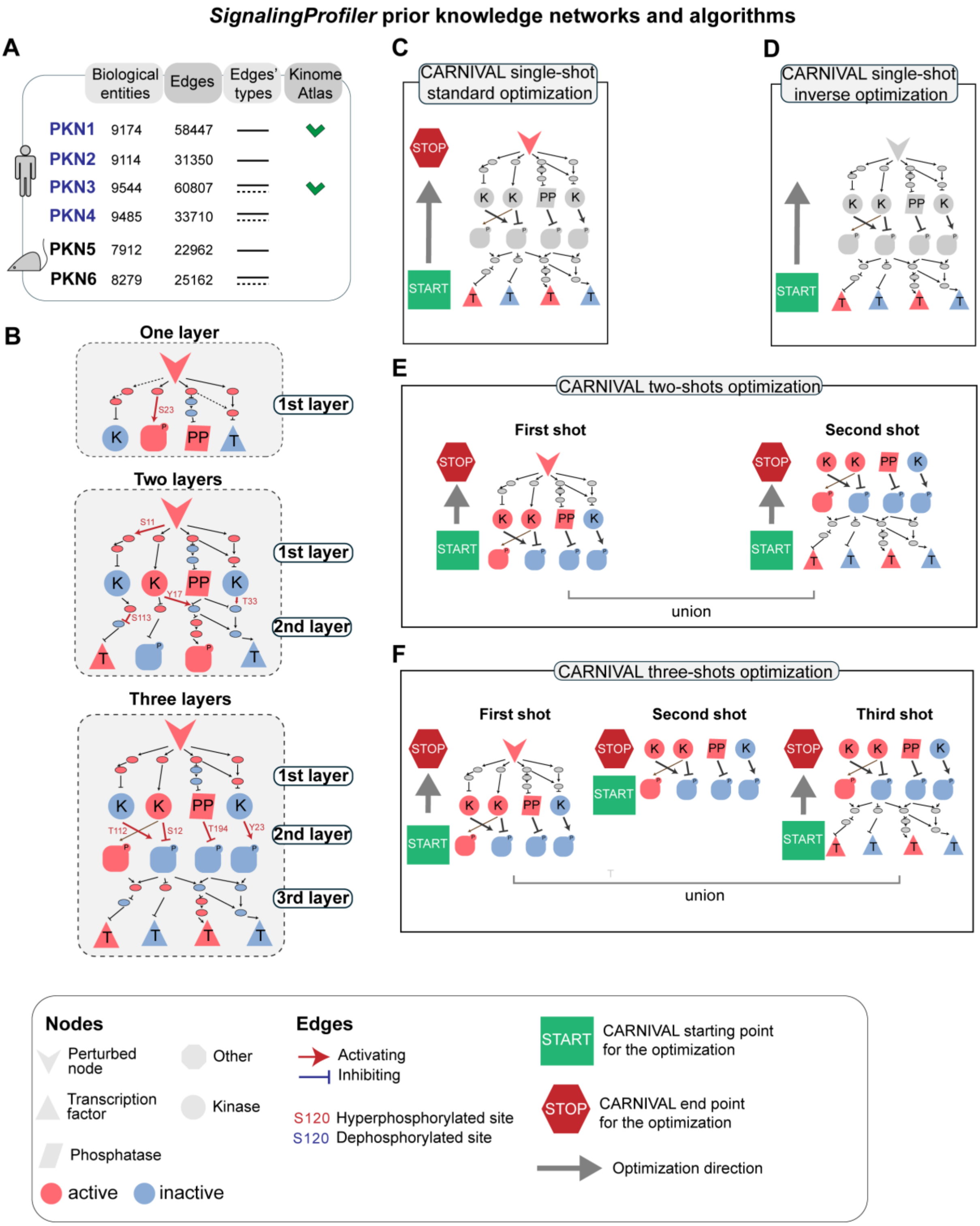


**Figure S3. *SignalingProfiler* prior knowledge networks (PKNs) and algorithms for network construction.**

**A.** Table representing the description and number of nodes and edges of the six prior knowledge networks (PKN).

**B.** Types of naïve networks that can be created by the shortest path algorithm. In the one-layer network, the perturbed node (starting point) is connected to all inferred proteins without differentiating their molecular functions. The two-layered network connects the perturbed node to kinases/phosphatases/others and then creates a second layer connecting them to transcription factors. The three-layered network adds another layer between kinases/phosphatases and other signaling proteins.

**C-F.** Different CARNIVAL implementations available in *SignalingProfiler*. The CARNIVAL algorithm, running in a single iteration, can be executed with (**C**) or without (**D**) a predefined endpoint (or perturbed nodes), denoted as the *standard* and *inverse* CARNIVAL, respectively. Additionally, two multi-shot versions are available in *SignalingProfiler.* In the two-step optimization, we first optimize from perturbed proteins to kinases, phosphatases, and other signaling proteins. This is followed by the optimization from the latter group to transcription factors, with subsequent union of the optimized sets (**E**). In the three-step optimization, the process involves consecutive runs: from perturbed proteins to kinases and phosphatases, then from kinases and phosphatases to other signaling proteins, and finally from the last set to transcription factors. Then, all the optimized submodels are combined (**F**).

#### Benchmarking of protein inference activity (Step 1) of *SignalingProfiler*

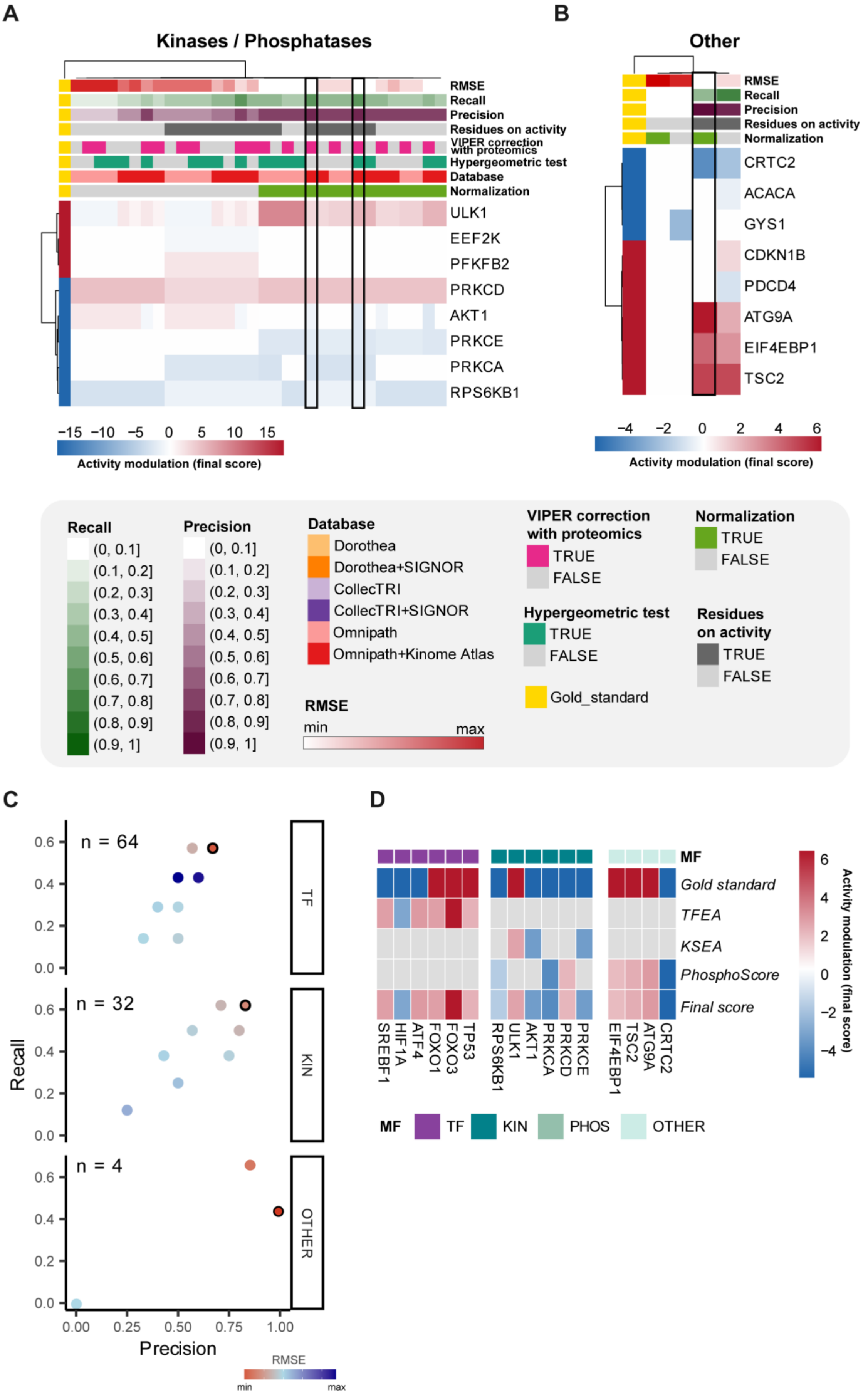

**Figure S4. Result of the benchmarking of *SignalingProfiler* protein inference (Step1).**

**A.** Heatmap reporting the inferred activity (*final score*) of gold standard kinases, using both footprint-based and PhosphoScore methods across 32 technical conditions (see Supplementary Material). Conditions encompass "Database" selection, correction with "Hypergeometric test" or proteomics ("VIPER correction with proteomics"), normalization of phosphoproteomics using proteomics ("Phospho normalization"), and the use of phosphosites regulating the sole activity ("Residues on activity"). Precision, recall, and Root Mean Squared Error (RMSE) are reported by comparing each condition to the gold standard (see Methods). The black box highlights the best technical conditions set as default in *SignalingProfiler*. Blue and red represent inactive and active proteins, respectively.

**B.** Heatmap reporting the inferred activity (*final score*) of other signaling proteins obtained by the PhosphoScore technique across 4 technical conditions, encompassing normalization of phosphoproteomics using proteomics ("Phospho normalization") and the use of phosphosites regulating the sole activity or also protein stability ("Residues on activity"). Precision, Recall, and Root Mean Squared Error (RMSE) are reported by comparing each condition to the gold standard (see Methods). The black box highlights the best technical condition set as default in *SignalingProfiler*. Blue and red represent inactive and active proteins, respectively.

**C.** Scatter plot of Precision (x-axis) and Recall (y-axis) for each technical condition in each molecular function. Dots are colored from red to blue according to increasing Root Mean Squared Error between the *final score* and the expected activity in the gold standard. The black-bordered dots are the best technical parameters selected as default for *SignalingProfiler*.

**D.** Heatmap summarizing the comparison between the expected activity ("Gold standard") and the activity score in the best technical condition of Step 1.

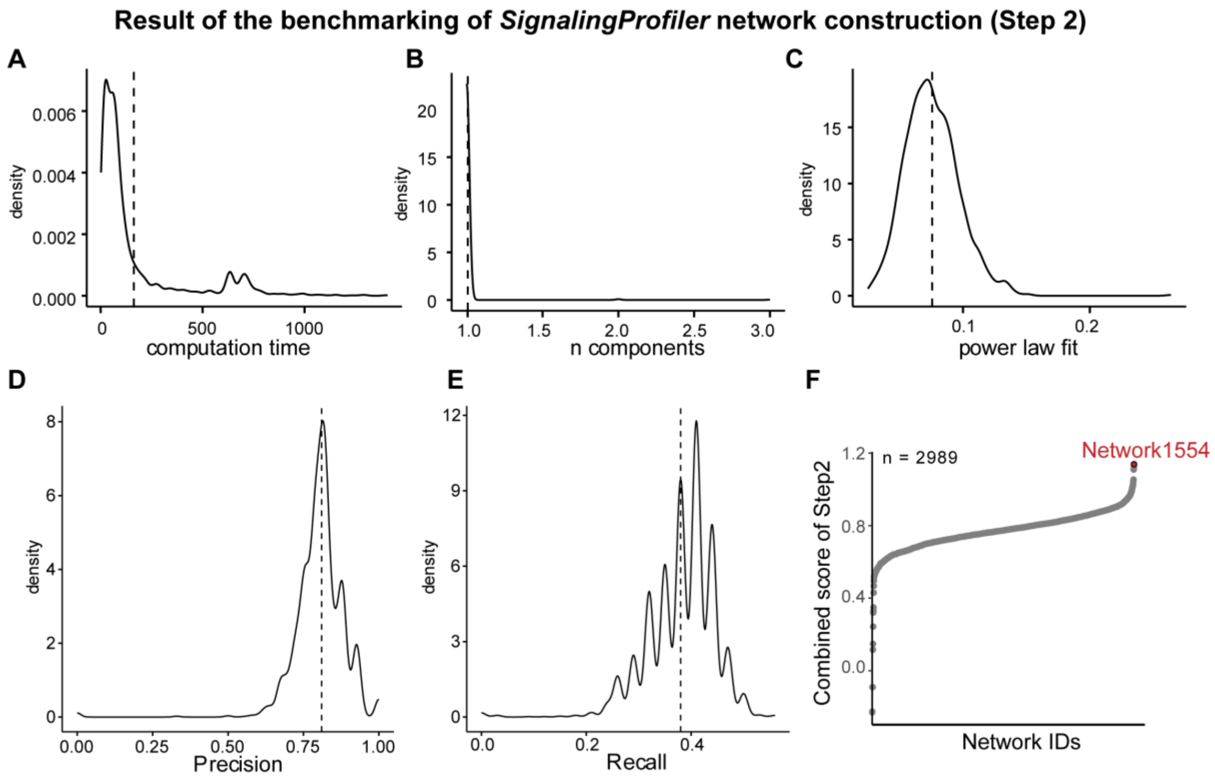


**Figure S5. Comparison of different network inference methods (Step 2).**

Distribution of different metrics across the family of 2989 generated models (Step 2) **(A-F):** computation time (**A**), number of components (**B**), fit to the power law (**C**), Precision (**D**), and recall (**E**) respect with the gold standard and the combined score of Step 2 exploited to rank the models (**F**).

**Benchmarking results of *SignalingProfiler* Step 2**

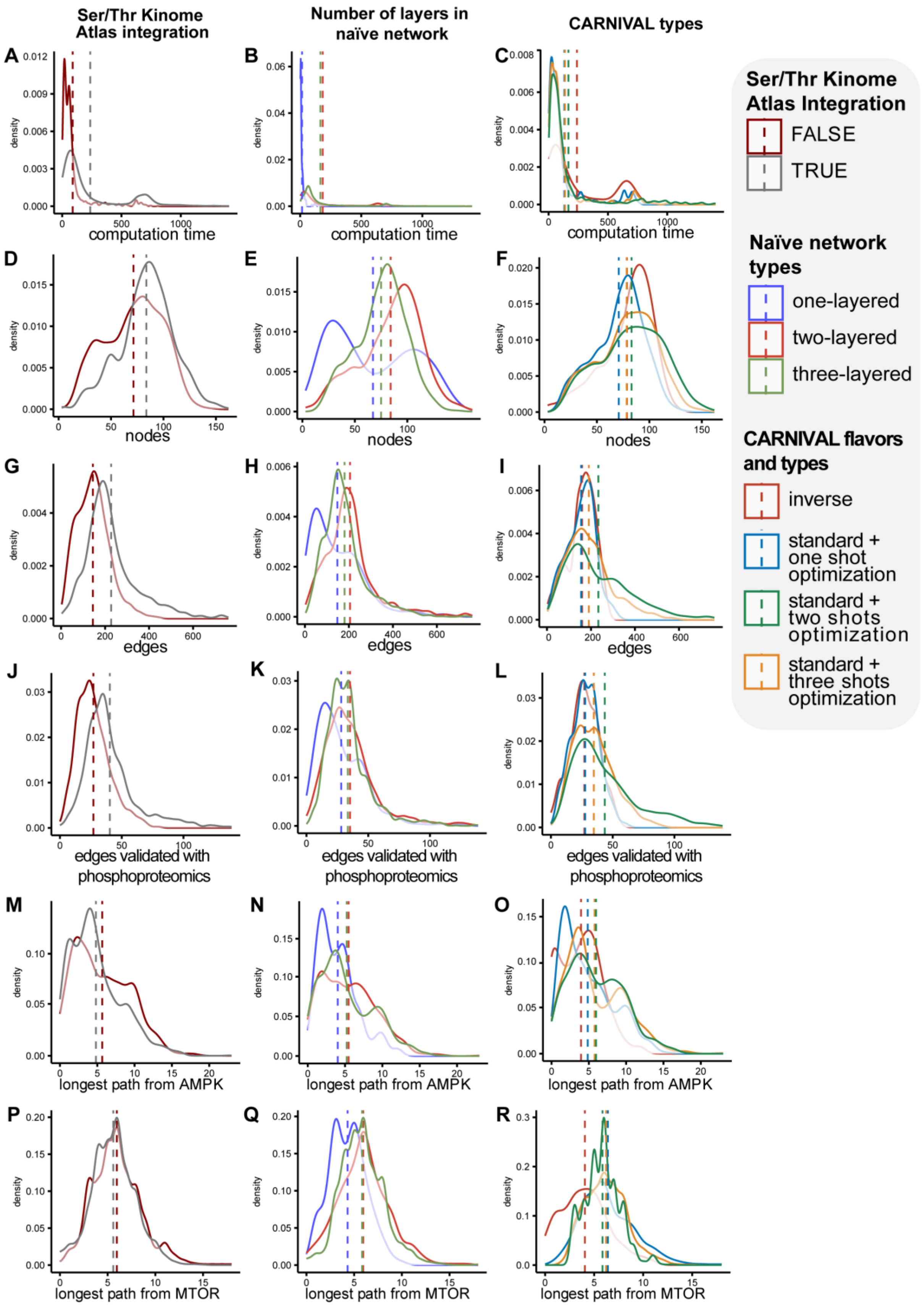

**Figure S6. Impact of Step 2 technical parameters on the resulting model properties.**

Impact of Ser/Thr Kinome Atlas integration (Johnson *et al*, 2023), number of layers in the naïve network, and CARNIVAL flavors and types on computation time (**A-C**), the number of nodes (**D-F**), edges (**G-I**), and of interactions validated by phosphoproteomics data (**J-L**), the maximum path length between end nodes (outdegree = 0) and AMPK (**M-O**) or mTOR (**P-R**), across 2989 generated models.

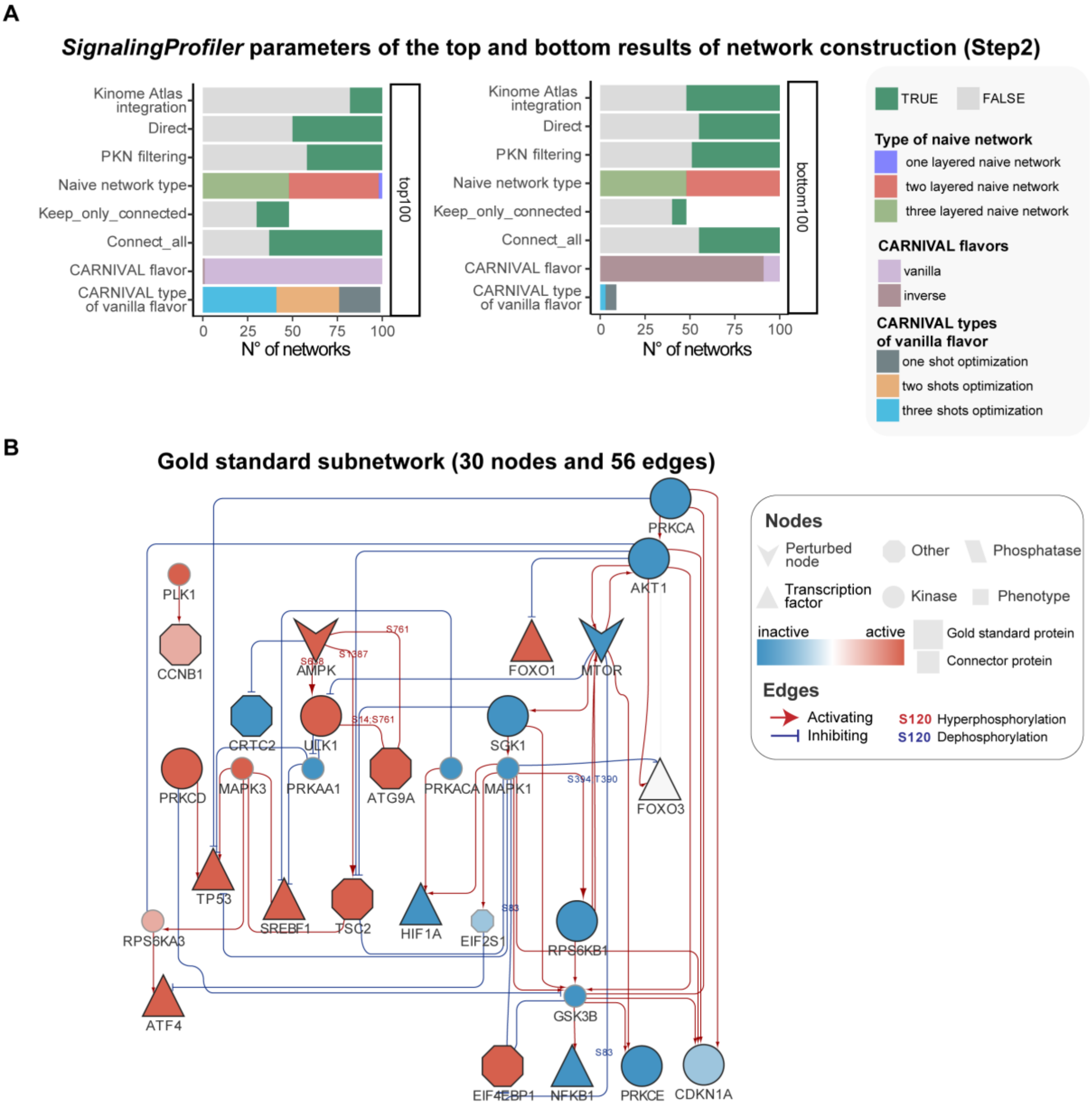


**Figure S7.** ***SignalingProfiler*** **top and bottom results of network construction (Step2) and gold standard subnetwork.**

**A.** Result of benchmarking of Step 2. Distribution of each parameter in the top and bottom 100 models after ranking 2989 produced models based on the combined score, considering adherence to the gold standard and topological features (see Methods). The most frequent parameters were chosen as default for Step 2.

**B.** Causal network representing a subnetwork extracted from the best *SignalingProfiler* model (**Figure S9**), with large and small nodes representing gold standard and connector proteins, respectively. Active and inactive proteins, along with activating and inhibitory edges, are indicated in red and blue, respectively. Phosphosites significantly modulated by metformin treatment and mapped onto edges are highlighted as edge labels.

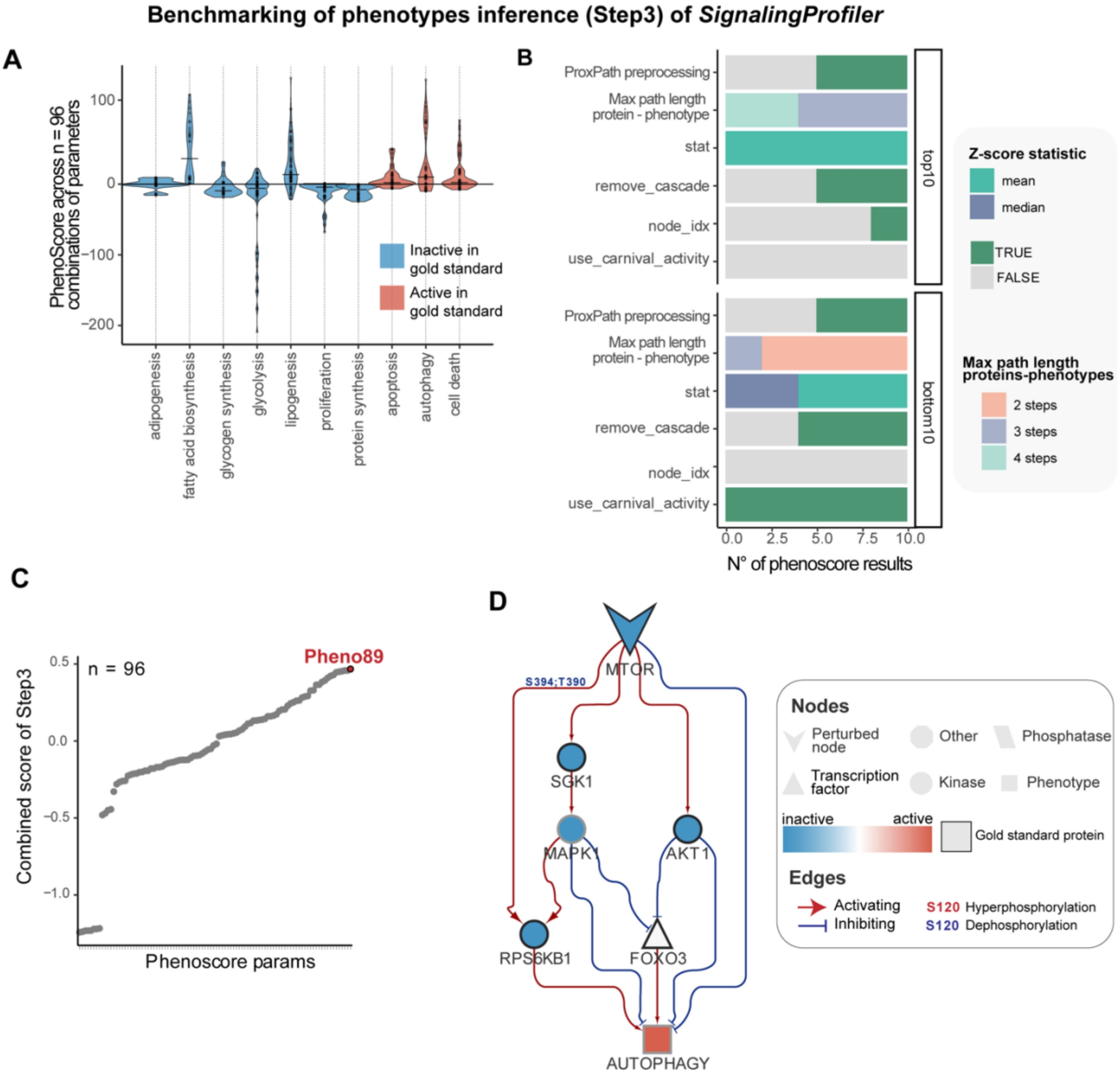


**Figure S8. Result of *SignalingProfiler* phenotype activity inference (Step 3) benchmarking.**

**A.** Violin plot illustrating the inferred activity distribution of 10 metformin-treatment-associated phenotypes across the 96 technical conditions of the benchmarking of *SignalingProfiler* Step 3. The color of each violin corresponds to the expected activity in the *phenotypic gold standard* (**Table S1**).

**B.** Result of benchmarking of Step 3. Distribution of each parameter in the top and bottom 10 phenotypes inference results after ranking 96 produced results based on the combined score that considers coherence with the phenotypic gold standard and computation time (see Methods).

**C.** Scatter plot displaying the distribution of the combined score across each of the 96 technical conditions evaluated to assess their fit to the *phenotypic gold standard* (see Methods). **D.** Functional circuit extracted from the *SignalingProfiler* best model linking mTOR inhibition to autophagy activation, as inferred from Step 3 of the pipeline.

#### *SignalingProfiler* resulting network (109 nodes and 309 edges)

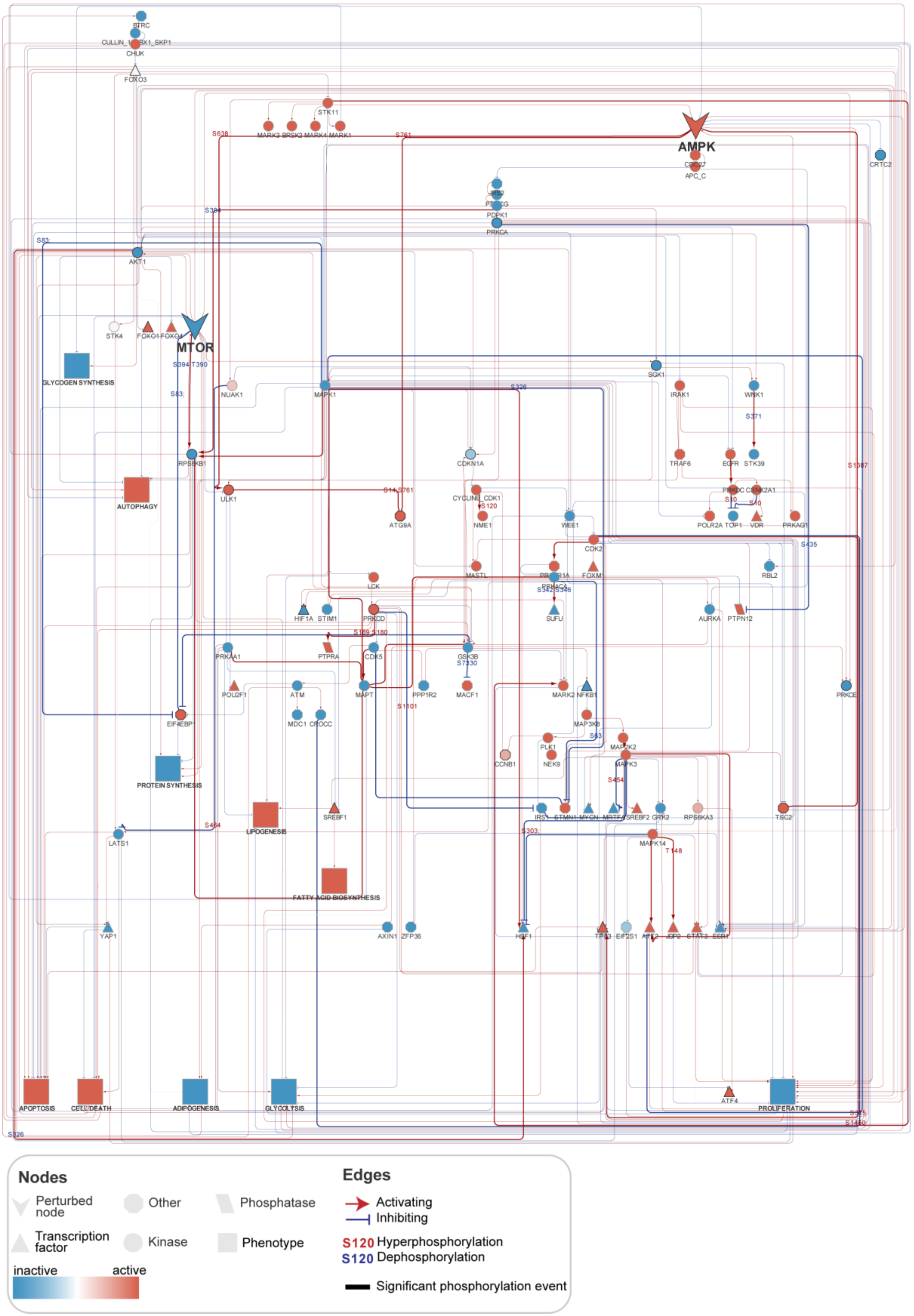

**Figure S9. The best result of the benchmarking process of *SignalingProfiler.*** Causal network of 109 nodes and 309 edges representing the metformin-induced signaling rewiring in breast cancer cells, starting from AMPK and mTOR proteins and ending on 10 relevant phenotypic traits. Activated and inhibited proteins after metformin treatment, along with activating and inhibiting edges, are indicated in red and blue, respectively. Node shape represents the molecular function. Phosphosites that are mapped on the interactions representing phosphorylation events are colored according to their level of phosphorylation after metformin treatment. Thicker edges represent (de)phosphorylations occurring at phosphosites significantly modulated in experimental data. Browse the model at https://www.ndexbio.org/viewer/networks/fa22e724-b54b-11ee-8a13-005056ae23aa.

#### Supplementary Tables Summary

**Table S1.** Protein and phenotypic gold standard table.

**Table S2.** Technical conditions for proteins' activity inference (Step 1) benchmarking.

**Table S3.** All results of proteins' activity inference (Step 1) benchmarking with quality metrics.

**Table S4.** Best result of proteins' activity inference (Step 1) benchmarking.

**Table S5.** Technical conditions and all results for network construction (Step 2) benchmarking.

**Table S6.** Technical conditions and all results for phenotypes' activity inference (Step 3) benchmarking.

**Table S7.** Best model of metformin-induced signaling rewiring returned by the whole benchmarking of *SignalingProfiler.*

#### Detailed explanation of *SignalingProfiler* functions and parameters

##### Step 1. Protein activity inference

*SignalingProfiler* extracts activity information for signaling proteins from experimental data through two methods: the footprint-based approach and PhosphoScore. If both methods generate an activity score for a protein, the scores are averaged; otherwise, only one of them is assigned. This process potentially results in the inference of 1174 transcription factors, 519 kinases, 62 phosphatases, and 1559 with other molecular functions.

*Footprint-based analysis*

Transcription factors-target genes collection was retrieved from Dorothea (confidence A) (Garcia-Alonso *et al*, 2019) and CollecTRI (Müller-Dott *et al*, 2023) resources using the *decoupleR* R package (Badia-I-Mompel *et al*, 2022) and from SIGNOR (filtering for transcriptional regulations) (Lo Surdo *et al*, 2023). Kinase-substrates and phosphatase-substrate collections were retrieved from Omnipath (Türei *et al*, 2016), using as sources only PhosphoSitePlus (Hornbeck *et al*, 2012), SIGNOR, HPRD (Goel *et al*, 2012), Reactome (Fabregat *et al*, 2018), phosphor.ELM (Dinkel *et al*, 2011), DEPOD (Damle & Köhn, 2019) databases. Kinase-substrate relations were integrated with Ser/Thr Kinome Atlas (Johnson *et al*, 2023) (see 'Ser/Thr Kinome Atlas parsing') and SIGNOR regulons updated to November 2023. To estimate transcription factors' activity from target genes and kinases and phosphatases from phosphosites, we used the VIPER algorithm (Alvarez *et al*, 2016). We used phosphosite/gene experimental fold change as statistics (VIPER parameters). We set the eset.filter parameter to FALSE. We included protein with at least 10 and 5 measured transcripts and phosphosites, respectively. We retained only proteins with enrichment p-value $< 0.05$. Inferred proteins that are not transcription factors or kinases/phosphatases according to the *SignalingProfiler* GO annotation algorithm (see 'Molecular function annotation' section) are discarded.

The hypergeometric test (using *phyper* from the *stats* package, version 4.1.2) was used to compute the likelihood of observing at least as many significant targets associated with a regulator in the dataset by random chance. A small p-value indicates an enrichment of significant targets linked to the regulator in the dataset. As such, we used the -Log10(p-value) to weight each protein VIPER score, giving more importance to proteins with a high number of significantly modulated targets.

To integrate multiple sources of evidence for a more robust interpretation, we correct VIPER output with proteomic. Specifically, when the VIPER output yields an insignificant fold-change that aligns with the direction of protein modulation, we correct this value to be significant. This refinement ensures effective integration when there is a consistent direction in the modulation of regulon targets and protein abundance.

*PhosphoScore*

To maximize the usage of phosphoproteomics data, we implemented a novel approach, dubbed *PhosphoScore,* to infer the activity of proteins undergoing de(phosphorylation) modifications. This approach integrates information regarding the regulatory role of phosphosites with their experimental fold-change. Leveraging data from SIGNOR and PhosphoSitePlus, we curated a table of regulatory phosphosites, distinguishing between those specifically affecting protein activity and those with broader effects (e.g., impacting the stability of the target protein). We excluded sites with ambiguous effects. Each phosphosite was identified by the primary gene name and a 15-mer centered on the modified residue. For both human and mouse datasets, a total of 6633 phosphosites were collected as regulators of activity or quantity (with 4325 exclusively affecting activity), associated with 2282 proteins. To enhance coverage in *Mus musculus*, instances where the regulatory role of mouse phosphopeptides was unknown were inferred from the human dataset utilizing the Blastp software (McGinnis & Madden, 2004).

To compute the activity of a phosphorylated protein we combined the regulatory role of its phosphosites with their fold-change in phosphoproteomics, as formulated by:

$$PhosphoScore = \frac{1}{n}\sum_{i=1}^{n} sign_i * FC_i$$

where: $n$ is the number of phosphosites regulating a protein, $sign_i$ is the regulatory role of phosphosite (1 or -1) and $FC_i$ is the experimental fold-change of phosphosites significantly modulated.

*Molecular function annotation*

We defined a simplified set of molecular functions for *SignalingProfiler*, namely transcription factors (‘tf’), kinases (‘kin’), phosphatases (‘phos’), and others (proteins not belonging to any of these categories). We created an annotated table that includes all the proteins found in the UniProt SwissProt proteome of both human and mouse using their GO: Molecular Function (MF) annotation and by querying the appropriate rest API of UniProt. Then, using the *getAncestors* function of the *GOSim* R package (v. 1.32) we retrieved the ancestors of each GO term. We assigned the molecular function using the GO term ('GO:0140110' for tf, 'GO:0016301' for kin, 'GO:0004725' for phos) or the ancestor GO term ('GO:0140110' for tf, 'GO:0016301' for kin, 'GO:0004725' for phos) associated to each protein. When a protein had an ambiguous annotation or was a protein complex, we manually annotated the molecular function.

**Step 2. Network construction**

*PKN preprocessing*

*SignalingProfiler* offers the possibility to remove from the PKNs the interactions between proteins not expressed in at least one omics layer between transcriptomic, proteomic, and phosphoproteomics.

*Naïve network generation*

*SignalingProfiler* restricts a user-defined PKN to the neighborhood of the inferred proteins in Step 1 and a set of user-defined perturbed nodes using the *shortest path algorithm.* The algorithm identifies all paths connecting two proteins within a maximum of 4 steps, terminating if a shorter path is found. Upon completion for all proteins, the *induce_subgraph* function of the *igraph* R package retrieves one-step interactions between proteins along the paths (**connect_all** parameter). Three frameworks were developed for creating neighborhoods with different layouts, determined by the number of layers.

A layer is defined by the connection using the shortest path algorithm of two types of molecular functions (see Molecular function annotation section). The *one-layered* network connects perturbed nodes to all inferred proteins without considering their molecular function. The *two-layered* network first connects the starting points to kinases/phosphatases/others and then creates a second layer from kinases/phosphatases/others to transcription factors by running the shortest path algorithm twice. The *three-layered* network introduces an extra layer between kinases/phosphatases and phosphoproteins running the shortest path algorithm with the max length parameter set to 1, aiming to select one-step direct interactions. Users can specify the max length of the shortest path for each run using the **max_length** parameter. Additionally, due to the independent execution of the shortest path algorithm in each layer, there may be cases where intermediate molecular functions like kinases, phosphatases, or phosphoproteins are connected within their layer but not to the layer above (for example, a kinase linked to its target but not to the perturbed node) or downstream (e.g., a phosphoprotein connected to its kinase but not to transcription factors). To address this, users can choose to eliminate proteins that are not connected to subsequent layers by utilizing the **keep_only_connected** parameter.

*Network optimization over inferred activity*

The naïve network undergoes optimization based on protein activity using the CARNIVAL R package (Liu *et al*, 2019). Through Integer Linear Programming (ILP), CARNIVAL selects edges in the naïve network whose signs align with the nodes' activity state (see (Liu *et al*, 2019; Dugourd *et al*, 2021) for details). The pipeline branches into Standard CARNIVAL (StdCARNIVAL) or Inverse CARNIVAL (InvCARNIVAL) based on the presence or absence of input nodes, respectively. The CARNIVAL algorithm aims to identify the smallest sign-coherent subnetwork and we exploited it to create four different optimization frameworks: two single-shot (**Figure S3C-D**) and two multi-shot frameworks (**Figure S3E-F**). The single-shot framework runs StdCARNIVAL and InvCARNIVAL algorithms in their original form. The multi-shot frameworks enhance model comprehensiveness. These frameworks independently optimize layers of the model, producing optimized subparts that are then combined.

Specifically, in the two-shot optimization process, StdCARNIVAL is first applied with the perturbed nodes as input and the kinases/phosphatases/others as output. Then, the second optimization shot focuses on optimizing connections specifically between kinases/phosphatases and others, connected in the former layer, to transcription factors. In the three-shot optimization, we introduce a third optimization step that specifically optimizes the kinases/phosphatases to other signaling proteins layer separately. The optimized graphs are then unified, and node activities in different subparts are averaged (**Figure S3C-F**).

**Step 3. Phenotypes activity inference and inclusion in the model (PhenoScore algorithm)**

The PhenoScore algorithm exploits and adapts the ProxPath algorithm (Iannuccelli *et al*, 2023) to infer the phenotypic traits' activity and incorporate them within the model. Briefly, ProxPath is a graph-based algorithm designed to measure the functional proximity of an input list of proteins to phenotypes, using causal interactions annotated in the SIGNOR database. Details are provided in (Iannuccelli *et al*, 2023).

*SignalingProfiler* provides a static table downloaded using ProxPath in November 2023 that accounts for 199 phenotypes. Moreover, *SignalingProfiler* can call ProxPath Python script using *reticulate* R package (v. 1.30-9000) to obtain an up-to-date table.

*Preprocess ProxPath phenotype-proximity table using experimental data*

The user can limit the ProxPath table to a subset of desired phenotypes (**desired_phenotypes** parameter). Moreover, in *SignalingProfiler*, users can contextualize the table, by excluding paths that involve proteins not expressed in at least one omics layer (transcriptomics, proteomics, or phosphoproteomics) (**preprocess** parameter).

*Run ProxPath algorithm*

The following operations of the ProxPath algorithm were translated in R and incorporated in the PhenoScore algorithm of *SignalingProfiler*.

Following a two-step process, ProxPath is employed to identify significantly proximal phenotypes. *Infer phenotype activity score (PhenoScore) from a user-defined protein list.*

The so-obtained table contains source proteins, target phenotypes, the proteins of each path (limited to 4 steps), and the p-value obtained from randomization. This table is merged with the node table of the *SignalingProfiler* optimized model, where each node is assigned an activity score.

The phenotype activity score is derived by computing the weighted mean of the activity of each phenotype regulator as follows:

$$PhenoScore = \frac{1}{n}\sum_{i=1}^{n} activity_i * sign_i * w$$

where the weight ($w$) is the -Log(p-value) obtained during the randomization process.

In cases where multiple proteins influence a common phenotype and one protein in the *SignalingProfiler* optimized model is downstream of another, users have the option to exclusively consider the most downstream protein as a phenotypic regulator (**'remove_cascade'** parameter).

To consider that a protein with a higher count of paths is more redundant and consequently a stronger regulator, we introduced the *redundancy* score (**node_idx**). For each phenotype, we count the proportion of paths originating from each protein over the total paths ending on that phenotype (**node_idx**). Optionally, the redundancy score can be multiplied and used as an additional weighting factor.

The activity of each phenotype regulator can be based on the CARNIVAL activity, available for all nodes in the model, or the final activity score, exclusive to the inferred proteins derived from the experimental data in Step 1 of the pipeline.

*Phenotypic circuits extraction*

The inferred phenotypes are integrated in the model adding an edge, representing an indirect interaction, from each regulator. *SignalingProfiler* provides a method to retrieve functional circuits starting from a user-defined set of nodes to a phenotype within a specified number of steps (k).
